# Golgi-dependent Reactivation and Regeneration of Quiescent Neural Stem Cells

**DOI:** 10.1101/2022.08.22.504877

**Authors:** Mahekta R. Gujar, Yang Gao, Xiang Teng, Qiannan Deng, Ye Sing Tan, Yusuke Toyama, Hongyan Wang

## Abstract

The ability of stem cells to switch between quiescent and proliferative states is crucial for maintaining tissue homeostasis and regeneration. In *Drosophila*, quiescent neural stem cells (qNSCs) extend a primary protrusion, which is removed prior to NSC reactivation. Here, we have unravelled that qNSC protrusions can be regenerated upon injury. This regeneration process relies on the Golgi apparatus which acts as the major acentrosomal microtubule-organizing centre in qNSCs. A Golgi-resident GTPase Arf1 and its guanine-nucleotide exchange factor Sec71 promote NSC reactivation and regeneration via the regulation of microtubule growth. Arf1 physically associates with its new effector Mini Spindles (Msps)/XMAP215, a microtubule polymerase. Finally, Arf1 functions upstream of Msps to target the cell-adhesion molecule E-cadherin to NSC-neuropil contact sites during NSC reactivation. Our findings have established *Drosophila* qNSCs as a new regeneration model and identified a novel Arf1/Sec71-Msps pathway in the regulation of microtubule growth and NSC reactivation.

## Introduction

The ability of stem cells to switch between quiescent and proliferative is crucial in maintaining tissue homeostasis. In the mammalian brain, the majority of adult neural stem cells (NSCs) exist in a quiescent state, without undergoing proliferation or differentiation ^1,2^. These quiescent NSCs (qNSCs) can be reactivated by various physiological stimuli such as injury or the presence of nutrition to take part in adult neurogenesis ^3^. Interestingly, the quiescent state of NSCs is associated with aging brains ^4^, and dysregulated NSC reactivation is associated with neurogenesis defects and neurodevelopmental disorders ^5,6^. Since current knowledge on NSC reactivation is extremely limited, more studies will be needed to gain a better understanding of the developmental, aging, and regeneration processes in the brain.

*Drosophila* larval brain NSCs or neuroblasts have emerged as a powerful model to understand the mechanisms underlying NSC quiescence and reactivation *in vivo* ^7,8^. In the *Drosophila* central nervous system, NSCs enter quiescence at the end of embryogenesis under the control of the spatial Hox protein, temporal identity factors, and the homeodomain differentiation factor Prospero. The qNSCs then resume proliferation mostly within 24h after larval hatching (ALH), in the presence of dietary amino acids ^9–11^. Dietary amino acids are sensed by the fat body, the functional equivalent of vertebrate liver and adipose tissue ^12,13^. The fat body generates mitogens, which presumably stimulate the glial cells forming the blood-brain barrier to secrete insulin-like peptides (dILPs). The dILPs promote reactivation of the underlying NSCs by locally activating the insulin receptor (InR)/phosphatidylinositol 3-kinase (PI3K)/Akt pathway in these cells ^14–16^. In mammalian brains, insulin-like growth factor-1 (IGF-1) and IGF-1 receptor (IGF-1R) have been shown to promote NSC proliferation ^17–19^. Furthermore, human IGF-1R mutations are associated with microcephaly, a neurodevelopmental disorder ^20^, suggesting that the insulin pathway is likely conserved from flies to humans and that similar mechanisms may be regulating NSC reactivation and proliferation in these organisms. Besides the InR pathway, other molecular machineries are known to regulate NSC reactivation. For instance, the spindle matrix complex intrinsically promotes NSC reactivation ^21^, while the Hippo pathway maintains NSC quiescence^3,7,8,10,22–27^

A hallmark of *Drosophila* qNSCs is their cellular protrusions, which extends from the cell body towards the neuropil and is removed prior to NSC reactivation ^15^. The biological functions of these protrusions are currently unclear. An earlier work from our lab has shown the presence of α-tubulin in the primary cellular extensions of qNSCs, suggesting that they are microtubule-enriched structures ^21^. Microtubules are polar filaments with a fast-growing plus-end and a slow-growing minus-end and have distinct orientations in the axons and dendrites of both invertebrate and vertebrate neurons. In *Drosophila*, they are oriented plus-end-out (plus ends distal to the cell body) in the axons and minus-end-out in the dendrites ^28^. Such distinct orientations are associated with different structural and functional properties of axons and dendrites, such as facilitating polarized neuronal cargo transport ^29^. In addition to facilitating cargo transport, the microtubules in *Drosophila* neurons also play an important role in axonal regeneration after injury, where destabilization enhanced dynamics, and reorganization of microtubule polarity are important for the regeneration of axons ^30^. In most dividing cells, including active NSCs, the centrosomes function as the microtubule-organizing centre (MTOC), the major site of microtubule nucleation and anchoring and the central point for the formation of microtubule arrays ^31,32^. Interestingly, a recent study from our lab has demonstrated that microtubules in the primary protrusion of qNSCs are predominantly acentrosomal and oriented plus-end-out, distal to the cell body ^33^. However, the microtubule-organizing centre of qNSCs is still unknown.

The Golgi apparatus and non-conventional Golgi structures such as Golgi elements and Golgi outposts have emerged as potential MTOCs in several cell types, including epithelial and muscle cells, and neurons ^34–36^. Golgi-derived microtubules have been shown to play important roles in post-Golgi trafficking, cell polarity establishment/maintenance, as well as neurite outgrowth and branching in neurons ^36^. ARF1, a member of the ADP-ribosylation factor (ARF) family of small G proteins, can recruit COPI coat proteins on *cis*-Golgi and clathrin adaptor proteins, such as AP-1, AP-3, and GGAs, on *trans*-Golgi in a GTP-dependent manner, thereby facilitating vesicle formation and trafficking ^37^. Mutations in critical Golgi proteins such as ARF1 and Sec71/ARFGEF2 (guanine-nucleotide exchange factor/GEF for ARF1) cause malformations of cortical development (MCD), a complex set of neurodevelopmental disorders in humans ^38,39^. However, how Arf1 and ARFGEF2/Sec71 contribute to brain development is unknown.

Mini spindles (Msps), an XMAP215/ch-TOG/Msps family protein, is a key regulator of microtubule growth in dividing cells ^40^; it functions as a microtubule polymerase that binds to microtubule plus ends ^40^. Our lab has previously demonstrated that Msps plays a vital role in NSC reactivation by regulating microtubule dynamics and orientation ^33^. We also showed that the cell adhesion molecule E-cadherin (E-cad), which localizes to NSC-neuropil contact sites in an Msps-dependent manner and with the help of the kinesin-2 complex (plus-end-directed microtubule motor proteins), is intrinsically required for NSC reactivation ^33^.

In this study, we have demonstrated for the first time that qNSC protrusions can be regenerated upon injury by laser severing. This regeneration relies on the protrusion’s proximal section, which we have shown to be enriched with Golgi apparatus and may thus function as the MTOC in qNSCs. We have identified critical roles for two Golgi proteins, Arf1 and its GEF Sec71, in NSC reactivation and regeneration via the regulation of microtubule growth. Furthermore, we have shown that Msps/XMAP215 functions downstream of Arf1 as its new effector, while, E-cad localizes to the NSC-neuropil contact sites, in an Arf1/Sec71-dependent manner, to promote reactivation of quiescent NSCs.

## Results

### *Drosophila* qNSC cellular protrusion can regenerate upon injury

To understand how *Drosophila* qNSC protrusions respond to injury, we examined the effect of laser ablation on the protrusions in *ex vivo* whole-mount larval brains (Figure 1A). qNSCs from larval brains at 0-2h ALH were ablated using a laser (Figure 1B, Panel 1, Movie S1, Methods), and subsequently imaged for up to 30 min using live imaging (Figure 1B, Panel 2-5, Movie S1). First, we severed the middle region of the qNSC protrusions, which is in between the cell body and the distal tip region. Remarkably, within 30 min of laser severing, most of the injured qNSC labelled by CD8-GFP (driven by *grainy head* (*grh*)-Gal4) completely repaired the injury on the protrusions (henceforth named regeneration) (Figure 1B, J, 5 in 6 qNSCs). Interestingly, we find that the distal end of the severed protrusion does not degenerate but is also capable of regeneration along with the proximal end (Figure 1B, Panel 1, Movie S1). The injured cellular protrusions were able to regenerate completely by 12 mins in 0-2h ALH *ex vivo* brains (Figure 1B). The average time taken to regenerate the protrusion following severing was 8.0 mins (Figure 1E). A minor case failed to fully regenerate within 30 min but showed partial regeneration, in which the proximal and distal ends of the severed protrusion had partial regrowth but fail to make final contact with each other within the 30 min imaging window (Figure 1J, 1 in 6 qNSCs).

**Figure 1.**
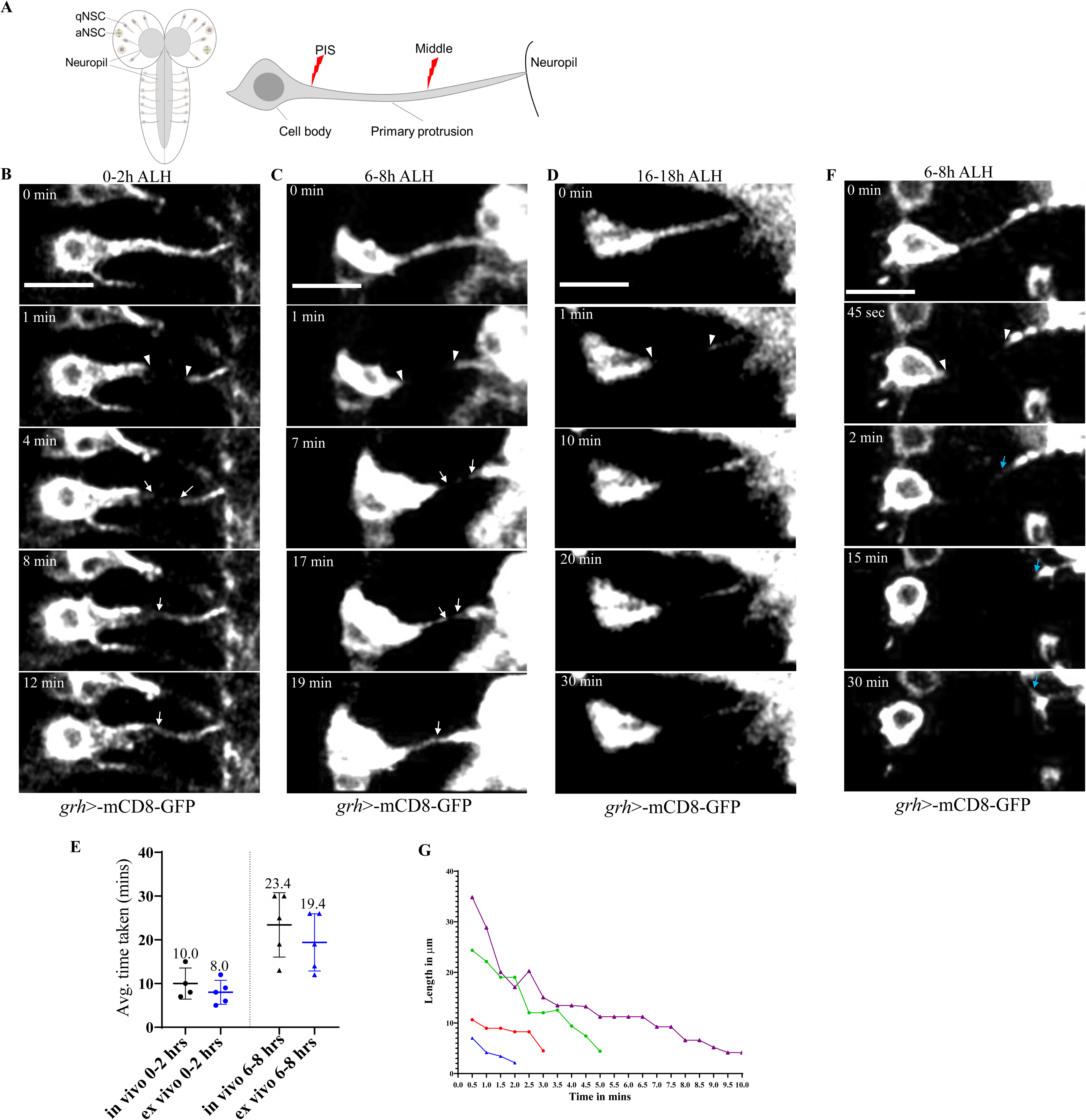
*Drosophila* quiescent NSC protrusions can regenerate upon injury. A) Illustrations of the *Drosophila* larval brain and laser ablation at different regions of qNSCs. PIS, Protrusion Initial Segment. B) Time series of a qNSC derived from *ex vivo* larval brain at 0-2h ALH from *grh*-Gal4; *UAS*-mCD8-GFP, ablated at the middle region of the protrusion (arrowheads). Complete regeneration is observed by 12 minutes. White arrows indicate regeneration of the protrusion after injury. Complete regeneration=83.3%, partial regeneration=16.7%, no regeneration=0% within 30 minutes of imaging. n=6. C) Time series of a qNSC derived from *ex vivo* larval brains at 6-8h ALH from *grh*-Gal4; *UAS*-mCD8-GFP, ablated at the middle region of the protrusion (arrowheads). Complete regeneration is observed by 19 minutes. White arrows indicate regeneration of the protrusion. Complete regeneration=62.5%, partial regeneration=25%, no regeneration=12.5% within 30 minutes of imaging. n=8. D) Time series of a qNSC derived from *ex vivo* larval brains at 16-18h ALH from *grh*-Gal4; *UAS*-mCD8-GFP, ablated at the middle region of the protrusion (arrowheads). Protrusion fails to regenerate. Complete regeneration=0%, partial regeneration=55.6%, no regeneration=44.4% within 30 minutes of imaging. n=9. E) Quantification of the time taken for complete regeneration of a qNSC with *grh*-Gal4; *UAS*-mCD8-GFP, in *in vivo* and *ex vivo* larval brains at 0-2h ALH and 6-8h ALH following laser ablation at the middle region of the protrusion. 0-2 h ALH: *in vivo*, 10 mins, n=4; *ex vivo*, 8 mins, n=5. 6-8 h ALH: *in vivo*, 23.4 mins, n=5; *ex vivo* 19.4 mins, n=5. F) Time series of a qNSC from *ex vivo* larval brains at 6-8h ALH from *grh*-Gal4; *UAS*-mCD8-GFP, ablated at the PIS (arrowheads). Blue arrows indicate retraction of the cellular protrusion attached to the neuropil region. 100% no regeneration, n=5. G) Quantification of the time taken for retraction of the cellular protrusion attached to the neuropil region, for qNSCs from larval brains at 6-8h ALH from *grh*-Gal4; *UAS*-mCD8-GFP when ablated at PIS region (G). Retraction was analyzed over a period of 30 minutes. n=5. Arrowheads indicate the ablated regions (B-D, G, I-J). White arrows indicate regeneration of the qNSC protrusions (B-C, I). Scale bars: 10 μm.

To validate our findings *in vivo*, we performed similar laser ablation experiments using intact larvae and severed the middle region of qNSC protrusions. Live imaging and ablation in whole larvae were performed using a PDMS microfluidic device, which immobilizes the animal using gentle mechanical force upon the application of vacuum ^41^. At 0-2h ALH, qNSC protrusions *in vivo* were also able to completely regenerate (Figure S1A; Movie S2). The average time taken to regenerate the protrusions following severing in 0-2h ALH *in vivo* brains was 10.0 mins (Figure 1E), suggesting that regeneration *in vivo* has a similar pace to that in *ex vivo* larval brains (8.0 min). The regeneration was not due to photobleaching and fluorescence recovery effect. Firstly, ablation of the cell body with the same laser power results in the death of the qNSCs within a couple of minutes (Figure S1B; Movie S3). Secondly, instantaneous recoil of the severed protrusion was observed in 60% of qNSCs (Figure S1A; n= 20; red arrows), demonstrating laser ablation, rather than photobleaching, on the protrusions. Thirdly, fluorescence recovery after photobleaching typically takes place within 1 minute after photobleaching, while in our experiments fluorescence re-appeared after 8 min or later following laser ablation. These observations present the first evidence that qNSC protrusions can be regenerated upon injury both *in vivo* and *ex vivo*.

### The regeneration of qNSC protrusion is developmentally- and spatially-regulated

Next, we investigated whether the regeneration of qNSC protrusions following injury depends on developmental stages or age. To this end, we severed the middle region of qNSC protrusions in *ex vivo* brains at different developmental stages and recorded the rate of regeneration. qNSC protrusions at 6-8h ALH were still capable of regenerating, although with reduced frequency compared to qNSCs at 0-2h ALH (Figure 1C, J; Movie S4). Furthermore, the average time for the complete regeneration of qNSC protrusions at 6-8h ALH was also significantly increased to 19.4 min compared to 8 min at 0-2h ALH (Figure 1E). Moreover, similar results were observed *in vivo* using whole larvae at 6-8h ALH (Figure 1E). Interestingly, at a later time point (16-18h ALH), none of the injured qNSC protrusions could regenerate completely; 55.6% of the qNSC protrusions showed partial regeneration while the remaining 44.4% showed no regeneration at all within 30 mins of imaging (Figure 1D, J; Movie S5). Therefore, our data demonstrated that the regeneration capability of qNSCs is developmentally regulated and declines rapidly over the first 18 hours of larval development. To understand the key factors in stage dependent regeneration capability, we sought to find a characteristic change in qNSCs over developmental stages. Acetylated-α-tubulin (Ace-tub on lysine 40), a posttranslational modification that stabilizes microtubules, is strongly enriched in the cell cortex as well as the cellular protrusions in qNSCs ^21^. We found that Ace-tub intensity in the protrusion gradually declines over the course of larval development (Figure S1C-F). Msps, an XMAP215/ch-TOG family protein, is a key regulator of microtubule growth in both quiescent and dividing NSCs ^33,42^, Interestingly, *msps* depletion led to significantly lower levels of Ace-Tub in the primary protrusions of qNSCs as compared to control brains at 0h and 6h ALH (Figure S1C-E), and almost similar levels at 16h ALH (Figure S1C and F). This data is consistent with age-dependent decline of the regeneration capability of qNSCs and suggests that the microtubules may play a role in the regeneration of qNSC primary protrusions.

To investigate the effect of the site of injury on the regeneration capability of qNSC protrusions, we ablated the ‘protrusion initial segment’ (PIS)—region demarcating the region between the soma and the cellular protrusion (Figure 1A). Interestingly, when the PIS of qNSCs in *ex vivo* larval brains at 6-8h ALH were ablated, 100% of qNSCs failed to regenerate and gradually lost their cellular protrusions over time (Figure 1F; Movie S6); the (distal) cellular protrusion attached to the neuropil region retracted within 30 minutes (Figure 1F-G). Furthermore, the cell bodies of the qNSCs changed shape from oval to a more rounded form over time, implying a change of cell morphology as a result of diminished mechanical tension following the loss of the entire protrusion. These data suggest that the regeneration of qNSC protrusions is spatially-regulated, with the PIS of the protrusions playing an indispensable role.

### The Golgi apparatus localizes to the protrusion initial segment and pericentrosomal regions in *Drosophila* qNSCs

Since our laser ablation experiments highlighted the importance of the PIS in the regeneration of qNSC protrusions, we sought to identify cellular organelles that are localized to the PIS region and responsible for the regeneration. Previous studies have shown the importance of Golgi in acentrosomal microtubule nucleation in neurons ^43^. In dividing NSCs, Golgi labelled with the *cis*-Golgi marker GM130 (100%, n=47) at 24h ALH were visualized as 5-15 small puncta per dividing NSC dispersed throughout the cytoplasm (Figure 2A, S2A). This was consistent with previously-reported dispersed localization pattern of the Golgi stacks in *Drosophila* cells ^44^. By contrast, distinct Golgi ribbons are located near the nucleus and the centrosome in most vertebrate cells ^45^. Since the Golgi distribution in qNSCs was unknown, we examined the localization pattern of Golgi-GFP in qNSCs at 6h ALH. Unexpectedly, trans-Golgi marker Golgi-GFP was observed as 1-3 large puncta predominantly localized to the PIS of qNSCs as well as to the putative pericentrosomal region at the apical side distal to the protrusions (Figure 2B and S2A; 100%, n=46). Occasionally, Golgi-GFP-positive puncta were observed in the cellular protrusions (Figure 2B; 4%, n=46). Furthermore, GM130 had similar localization patterns in qNSCs as that of Golgi-GFP: only 1-3 GM130-positive large puncta were observed in qNSCs, all of which were located at the PIS and the apical region (Figure 2C). We observed an identical localization pattern of GM130 in qNSCs at 0h ALH (Figure 2D; 100%, n=26). In addition, medial-Golgi marker α-MannosidaseII (ManII-Venus) showed similar localization patterns in qNSCs at 6h ALH (Figure S2B). Interestingly, the size of the Golgi puncta in qNSC was temporally-regulated; the size of Golgi inferred by GM130 was significantly smaller at 16h ALH than 6h ALH, with a further reduction in dividing NSCs at 24h ALH (Figure 2A, C, E-F). These observations indicate that the Golgi has a unique localization at the PIS in qNSCs, distinct from that in dividing NSCs.

**Figure 2.**
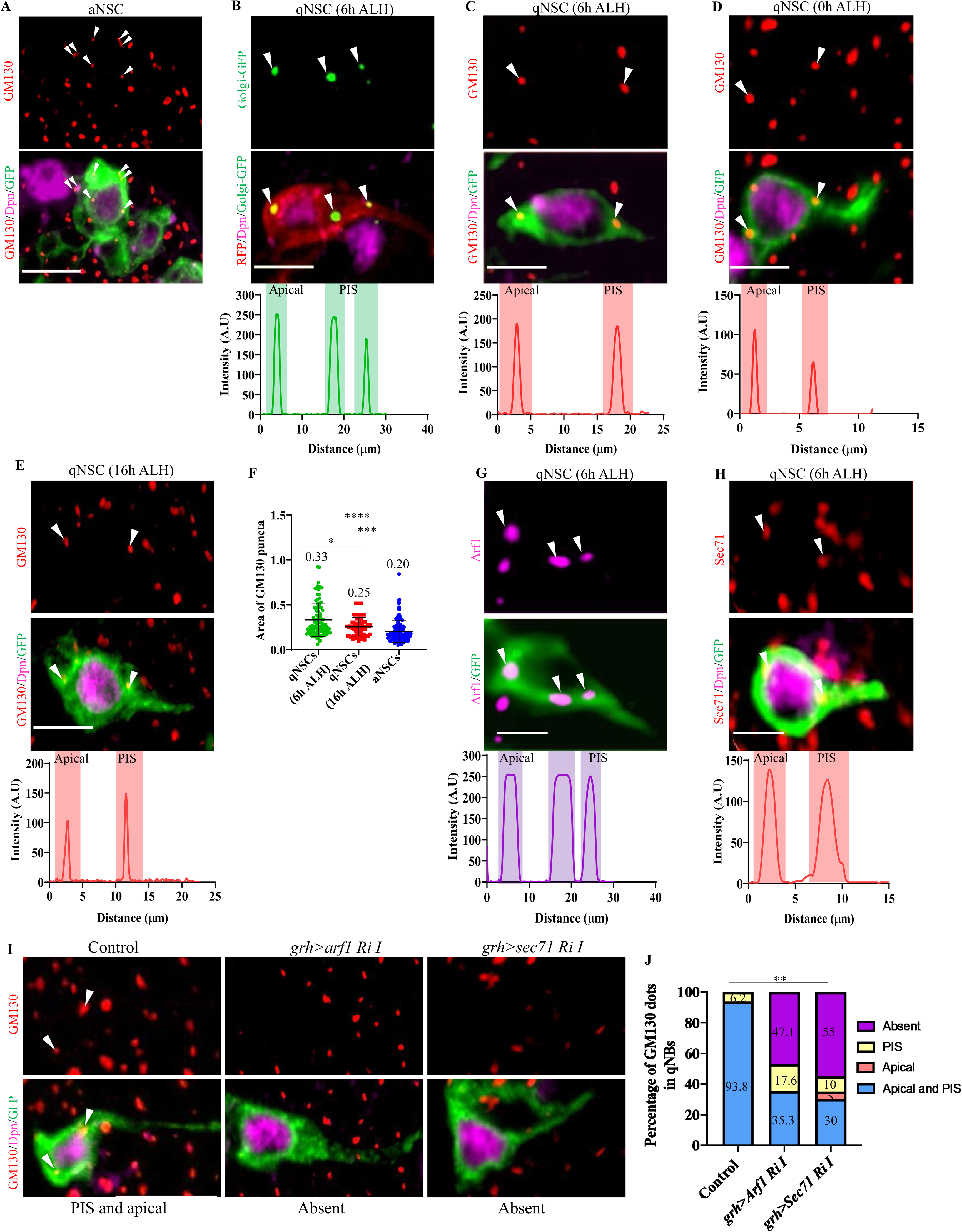
The Golgi apparatus localizes to the protrusion initial segment in *Drosophila* qNSCs. A) A dividing NSC in the larval brain expressing *grh*>CD8-GFP at 24h ALH was labelled with antibodies against Dpn, GM130, and GFP.GM130 puncta located randomly in the cytosol, n=47. Arrowheads indicate GM130 localization. B) qNSCs in the larval brain expressing Golgi-GFP and CD8-RFP driven by *grh*-Gal4 at 6h ALH were labelled with antibodies against Dpn, GFP, and RFP. Arrowheads indicate Golgi-GFP puncta localization at the apical and PIS regions (100%, n=46). Intensity graphs showing the localization of Golgi-GFP in the apical and PIS regions of qNSCs. C) A qNSC in the larval brain expressing *grh*>CD8-GFP at 6h ALH was labelled with antibodies against Dpn, GM130, and GFP.GM130 puncta located at apical and PIS, 100%, n=51. Arrowheads indicate GM130 puncta. Intensity graphs show localization of GM130 in the apical and PIS regions of qNSCs. D) A qNSC in the larval brain expressing *grh*>CD8-GFP at 0h ALH was labelled with antibodies against Dpn, GM130, and GFP. GM130 puncta located at apical and PIS, 100%, n=24. Arrowheads indicate GM130 localization. Intensity graphs show localization of GM130 in the apical and PIS regions of qNSCs. E) A qNSC in the larval brain expressing *grh*>CD8-GFP at 16h ALH was labelled with antibodies against Dpn, GM130, and GFP. GM130 puncta located at apical and PIS, 91.7%, n=36. Arrowheads indicate GM130 localization. F) Quantification graph of GM130 puncta areas in genotypes from (A-C). The average area of GM130 puncta in: qNSCs at 6h ALH=0.33µm^2^, n=101; qNSCs at 16h ALH=0.25 µm^2^, n=84; and aNSCs at 24h ALH=0.20 µm^2^, n=132. G) A qNSC protrusion in the larval brain expressing *grh*>CD8-GFP at 6h ALH was labelled with antibodies against Arf1 and GFP. Arrowheads indicate Arf1 puncta at both apical and PIS=100% n=45. Intensity graphs show Arf1 localization in the apical, PIS and lateral regions of qNSCs. H) A qNSC protrusion in the larval brain expressing *grh*>CD8-GFP at 6h ALH was labelled with antibodies against Sec71, Dpn, and GFP. Sec71 puncta at PIS, 95%; apical, 93%, n=31. Arrowheads indicate Sec71 puncta. Intensity graphs show Sec71 localization at the apical and PIS regions of qNSCs. I) qNSC protrusions in the larval brain at 6h ALH from control (*grh*-Gal4; *UAS* Dcr2/*UAS-β-Gal* RNAi), *arf1* RNAi I (*UAS*-*arf1* RNAi (VDRC#23082GD)) and *sec71* RNAi I (*UAS*-*sec71* RNAi (VDRC#100300KK)) expressing *grh*-Gal4 CD8-GFP were labelled with antibodies against GM130, Dpn and GFP. Arrowheads indicate GM130 localization. J) Quantification graph of the percentage of GM130 puncta per qNSC for genotypes in (I). GM130 puncta localization patterns: control, apical + PIS=93.8%, PIS only =6.2% (n=21); *arf1* RNAi I, apical + PIS=35.3%, PIS only =17.6%, absent =47.1% (n=29); *sec71* RNAi I, apical + PIS=30%, apical only =5%, PIS only =10%, absent =55% (n=32). **p<0.01. Scale bars: 5 μm.

### Arf1 and its GEF Sec71/ARFGEF2 localize to the protrusion initial segment and pericentrosomal region in qNSCs

We examined the localization of the Golgi protein Arf1 in qNSCs at 6h ALH. Similar to Golgi-GFP and GM130, Arf1 was always observed as 1-3 large puncta located at the PIS and the apical region of wild-type qNSCs (Figure 2G); Arf1 was also observed at the lateral region of the cell body in 5% (Figure 2G; n=45) of qNSCs. Likewise, intense localization of Arf1 was detected at the PIS and apical regions in qNSCs overexpressing Arf1-GFP at 6h ALH (Figure S2C).

Arf1 can be activated from a GDP-bound state to a GTP-bound state by Sec7 domain-containing GEFs ^46^. *Drosophila* Sec71 is one such GEF that plays an important role in regulating Arf1 functions during neuronal development ^47^. We, therefore, examined the localization pattern of Sec71 in qNSCs using anti-Sec71 antibodies. Similar to Arf1, at 6h ALH, Sec71 localized primarily to the PIS region, while Sec71-positive puncta were also observed at the apical region of qNSCs (Figure 2H). Taken together, the Golgi proteins Arf1 and Sec71 predominantly localize to the PIS region of qNSCs, from where the primary protrusions are extended.

Knockdown of *arf1* or *sec71* by RNAi caused a significant reduction in GM130 puncta in qNSCs, compared with the control at 6h ALH (Figure 2I-J). Similarly, upon overexpression of Sec71^E794K^, a dominant-negative form of Sec71 (hereafter referred to as Sec71^DN^) at 6h ALH, GM130 puncta were missing from the PIS region in 56% of qNSCs (Figure S2D-E). These data suggest that Arf1 and its GEF Sec71 are required for the proper localization of GM130 in qNSCs.

Although the Golgi punctum was seen at the putative pericentrosomal region at the apical side in qNSCs (Figure 2B), Golgi marked by GM130 and the immature centrosome marked by a constitutive centriolar protein Asterless (Asl) localize “near to each other” (0.5-1.5 microns distance), without any overlap, in most qNSCs at 6h ALH (Figure S2F 63.9%). Golgi is located far apart from the centrosome (>1.5 microns distance) in 22.2% qNSCs, while GM130 partially colocalized with Asl (<0.5 microns) in 13.9% of qNSCs (Figure S2G-H), but complete co-localization was never seen in wild-type qNSCs. Similar localization patterns were observed with Arf1 and Sas-4, another centriolar protein in majority of qNSCs at 6h ALH (Figure S2I-K). Sec71 co-localized with Arf1 and partially co-localized with GM130 at the PIS, apical or lateral regions in qNSCs (Figure S2L-M). Similar to Arf1, Sec71 mostly localized close to Asl, and occasionally localized either far away or partially overlapping with Asl (Figure S2N-P). These data indicate that the apical Golgi localizes to pericentrosomal region but rarely overlaps with the immature centrosome in qNSCs.

### The Golgi proteins Arf1 and its GEF Sec71 are required for qNSC reactivation

The mechanism by which Arf1 and its GEF ARFGEF2/Sec71 contribute to brain development is unknown. To understand the functions of Arf1 in neurodevelopment, we first examined *arf1* loss-of-function phenotypes during NSC reactivation. To do this, we fed the larvae with EdU-containing food for 4h so that all cycling NSCs were incorporated with EdU as described previously (Li et al., 2017). Interestingly, at 24h ALH, *arf1* knockdown using two independent RNAi lines showed dramatic increase in EdU-negative NSCs compared with the control (Figure 3A-B). Moreover, there was a significant increase in the percentage of qNSCs retaining the primary protrusion upon *arf1* knockdown, compared with control (Figure 3C). Furthermore, the delayed NSC reactivation observed in EdU incorporation experiments upon *arf1* RNAi were fully rescued at 48h ALH and partially rescued at 24h ALH by the expression of an RNAi-resistant *arf1* transgene (Figure 3D-F), confirming the specificity of the *arf1* RNAi effect. The better rescue at 48h ALH is likely due to higher expression levels of the RNAi-resistant Arf1 transgene as compared to 24h ALH (Figure S3A-D).

**Figure 3.**
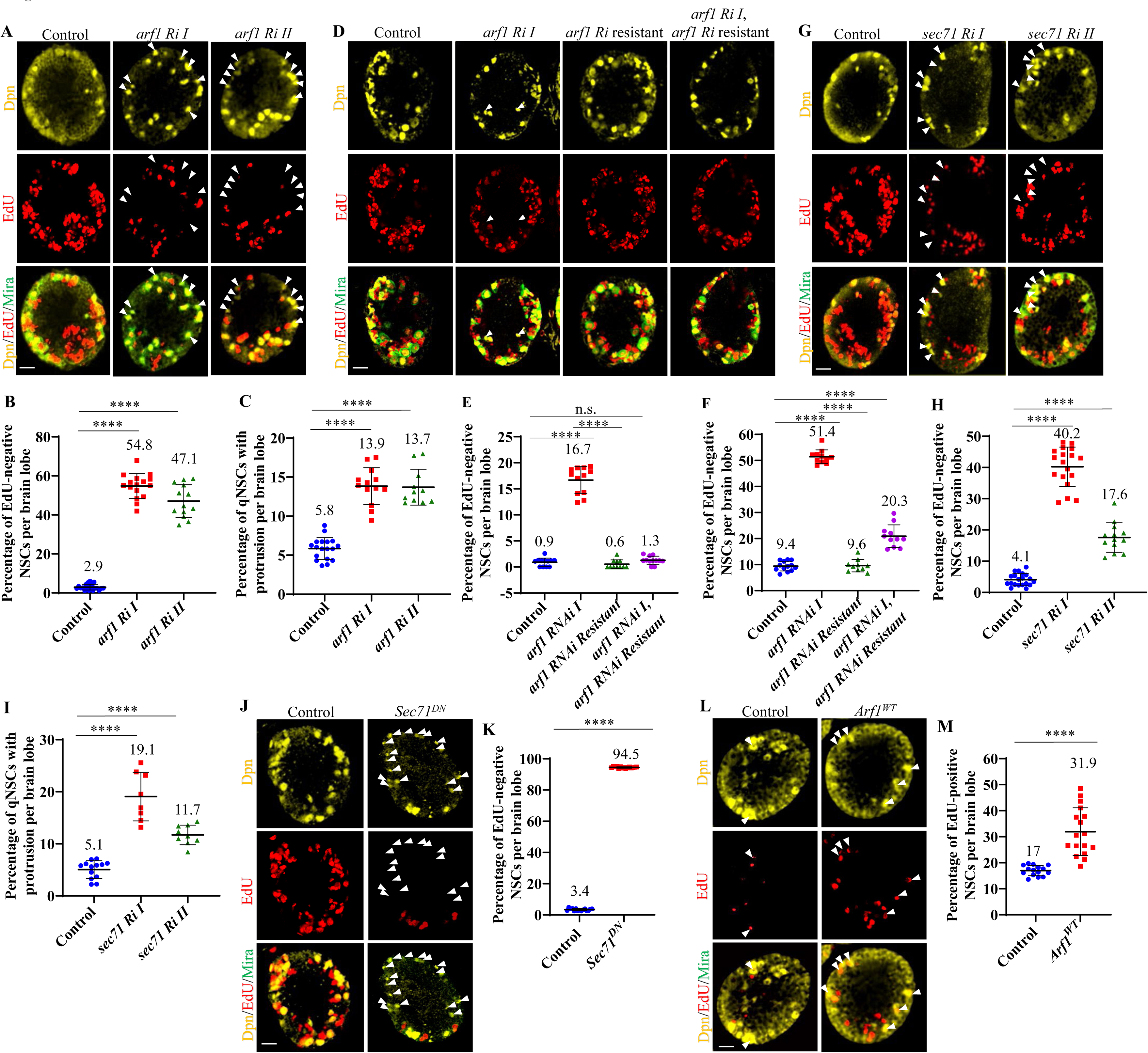
The Golgi proteins Arf1 and its GEF Sec71 are required for qNSC reactivation. A) Larval brains at 24h ALH, including control (*grh*-Gal4; *UAS*-dicer2 *UAS*-*β-Gal* RNAi), *arf1* RNAi I (*UAS*-*arf1* RNAi /VDRC#23082GD) and *arf1*RNAi II (*UAS*-*arf1* RNAi/VDRC#103572KK), driven by *grh*-Gal4; *UAS*-*Dicer2* were analysed for EdU incorporation and labelled with Dpn and Mira. B) Quantification graph of percentage of EdU-negative NSCs per brain lobe for genotypes in (A). Control, 7.6%, n=20 brain lobes (BL); *arf1 RNAi* I, 54.8%, n=16 BL; *arf1* RNAi II, 47.1%, n=13 BL. ****p<0.0001. C) Quantification graph of percentage of qNSCs with protrusions per brain lobe for genotypes in (A). Control, 5.8%, n=18 BL; *arf1* RNAi I, 13.9%, n=13 BL; *arf1* RNAi II, 13.7%, n=11 BL. ****p<0.0001. D) Larval brains at 48h ALH, including the control (*grh*-Gal4 *UAS-β-Gal* RNAi), *UAS*-*arf1 RNAi* I, *UAS*-*arf1 RNAi* I resistant, and *UAS*-*arf1 RNAi* I, *UAS*-*arf1 RNAi* I resistant, driven by *grh*-Gal4 were analyzed for EdU incorporation and labelled with Dpn and Mira. E) Quantification graph of percentage of EdU-negative NSCs per brain lobe for genotypes in (D). Control, 0.9% n=12 BL; *arf1 RNAi* I, 16.7% n=12 BL; *arf1* RNAi I resistant, 0.6%, n=11 BL; *arf1* RNAi I, *arf1* RNAi I resistant 1.3%, n=12 BL. ****p<0.0001, ns-non-significant. F) Quantification graph of percentage of EdU-negative NSCs per brain lobe for genotypes at 24h ALH, including the control (*grh*-Gal4 /*UAS-β-Gal* RNAi), *UAS*-*arf1 RNAi* I, *UAS*-*arf1 RNAi* I resistant, and *UAS*-*arf1 RNAi* I, *UAS*-*arf1 RNAi* I resistant, driven by *grh*-Gal4. Control, 9.4% n=13 BL; *arf1 RNAi* I, 51.6% n=11 BL; *arf1* RNAi I resistant 9.6%, n=10 BL; *arf1* RNAi I, *arf1* RNAi I resistant 20.3%, n=11 BL. ****p<0.0001, ns-non-significant. G) Larval brains at 24h ALH, including the control (*grh*-Gal4, *UAS*-dicer2 *UAS*-*β-Gal* RNAi), *sec71* RNAi I (*UAS*-*sec71* RNAi /VDRC#100300KK) and *sec71* RNAi II (*UAS*-*sec71* RNAi/BSDC#32366), driven by *grh*-Gal4; *UAS*-Dicer2 were analyzed for EdU incorporation and labelled with Dpn and Mira. H) Quantification graph of percentage of EdU-negative NSCs per brain lobe for genotypes in (G). Control, 4.1% n=20 BL; *sec71* RNAi I, 40.22% n=18 BL; *sec71* RNAi II, 17.6% n=13 BL. ****p<0.0001. I) Quantification graph of percentage of qNSCs with protrusions per brain lobe for genotypes in (G). Control, 5.1%, n=13 BL; *sec71* RNAi I, 19.1%, n=8 BL; *sec71* RNAi II, 11.7%, n=9 BL. ****p<0.0001. J) Larval brains at 24h ALH, including the control (*grh*-Gal4/*UAS*-*β-Gal* RNAi) and *UAS*-*Sec71^DN^*, driven by *grh*-Gal4 were analyzed for EdU incorporation and labelled with Dpn and Mira. K) Quantification graph of percentage of EdU-negative NSCs per brain lobe for genotypes in (J). Control, 3.4%, n=15 BL; *UAS*-*Sec71^DN^*, 94.5%, n=15. ****p<0.0001. L) Larval brains at 6h ALH, including the control (*grh*-Gal4 *UAS*-*β-Gal* RNAi) and Arf1 overexpression (*UAS*-*Arf1^WT^*), driven by *grh*-Gal4 were analyzed for EdU incorporation and labelled with Dpn and Mira. Arrowheads indicate EdU-positive NSCs. M) Quantification graph of percentage of EdU-positive NSCs per brain lobe for genotypes in (L). Control, 17%, n=15 BL; *UAS*-*Arf1^WT^*, 31.9%; n=17 BL. ****p<0.0001. Arrowheads indicate EdU-negative NSCs (A, D, G, J) or EdU-positive NSCs (L). Scale bars: 10 μm.

Next, we investigated whether Sec71 is required for NSC reactivation. The percentage of EdU-negative qNSCs was dramatically increased upon *sec71* knockdown by two independent RNAi lines compared with the control at 24h ALH (Figure 3G-H), suggesting a significant delay in NSC reactivation. The percentage of qNSCs retaining the primary protrusion in *sec71* knockdown larval brains was also significantly increased as compared to control brains (Figure 3I). At 24h ALH, the reactivation defects caused by *sec71* knockdown, as observed in EdU incorporation experiments, was well-rescued by expressing a RNAi-resistant *sec71* transgene (Figure S3E-F). *Sec71^DN^* overexpression in NSCs also caused a severe defect in NSC reactivation; 94.5% of *Sec71^DN^* NSCs failed to incorporate EdU as compared to 3.4% of wild-type NSCs (Figure 3J-K).

Taken together, our results indicate that the Golgi proteins Arf1 and Sec71 are key regulators of NSC reactivation.

### Overexpression of *arf1* triggers premature NSC reactivation

Since the loss of *arf1* and *sec71* led to delayed reactivation of qNSCs, we wondered whether overexpressing Arf1 or Sec71 had any effect on NSC reactivation. At 6h ALH, most of the wild-type NSCs were still in a quiescent state, with about 17% of NSCs incorporating EdU (Figure 3I-J). Interestingly, the overexpression of *Arf1* (*UAS*-*Arf1^WT^*) driven by *grh*-Gal4 resulted in significantly more proliferative NSCs (Figure 3L-M, 31.9%), suggesting that qNSCs were reactivated prematurely. However, in the absence of dietary amino acids (sucrose-only food), *Arf1* overexpression did not have any effect on reactivation (Figure S3G-H). In contrast, overexpression of *Sec71* (*Sec71^WT^*) did not cause premature NSC reactivation (15.7% EdU-positive NSCs, n=6 brain lobes (BLs)) as compared to the control at 6h ALH (12.8%, n=6 BL). These results indicate that Arf1 is both necessary and sufficient for NSC reactivation in the presence of nutrition.

### COPI and AP-1 are not important for NSC reactivation

Arf1 recruits coatomer complex proteins (COPI) and clathrin adaptor proteins (AP-1) at cis- and trans-Golgi network respectively during membrane trafficking ^48,49^. We tested whether COPI coatomer complex and AP-1 could regulate reactivation of qNSCs. Interestingly, knocking down of various AP-1 or COP1 subunits have no or subtle effects on NSC reactivation (Figure S3I-K), suggesting that the function of Arf1 in promoting NSC reactivation is likely largely independent of its well-known role in membrane trafficking.

### Acentrosomal microtubule growth occurs at the Golgi in qNSCs

To investigate whether the Golgi apparatus could be the MTOC in qNSCs, we examined whether microtubule growth can occur at the vicinity of Golgi in qNSCs. Microtubule growth was analyzed by tracking the movement of the End binding 1 (EB1)-GFP, a plus-end tracking protein (+TIP) that binds to microtubule plus-ends during microtubule growth. The position of emerging EB1-GFP comets can be used as a proxy for microtubule nucleation sites ^50,51^. The Golgi was marked by Golgi-GFP in these qNSCs. At 6h ALH, 65% of microtubules originated from the vicinity of Golgi at the PIS region of qNSCs (Figure 4A-B, 228 out 353 comets, Movie S7), whereas the remaining 35% of microtubules originated from the Golgi located at the apical or lateral regions (Figure S4A, Movie S8). Golgi-GFP did not interfere with our analysis of EB1-GFP movement, as Golgi-GFP puncta are stationary in live imaging and are much larger in size than the EB1-GFP comets. Besides, Golgi-GFP overexpression did not alter the velocity or direction of EB1 comets in qNSCs, although there was a minor but insignificant increase in the number of comets (Figure S4B-E). Therefore, acentrosomal microtubules emerge mostly from the Golgi at the PIS region in qNSCs.

**Figure 4.**
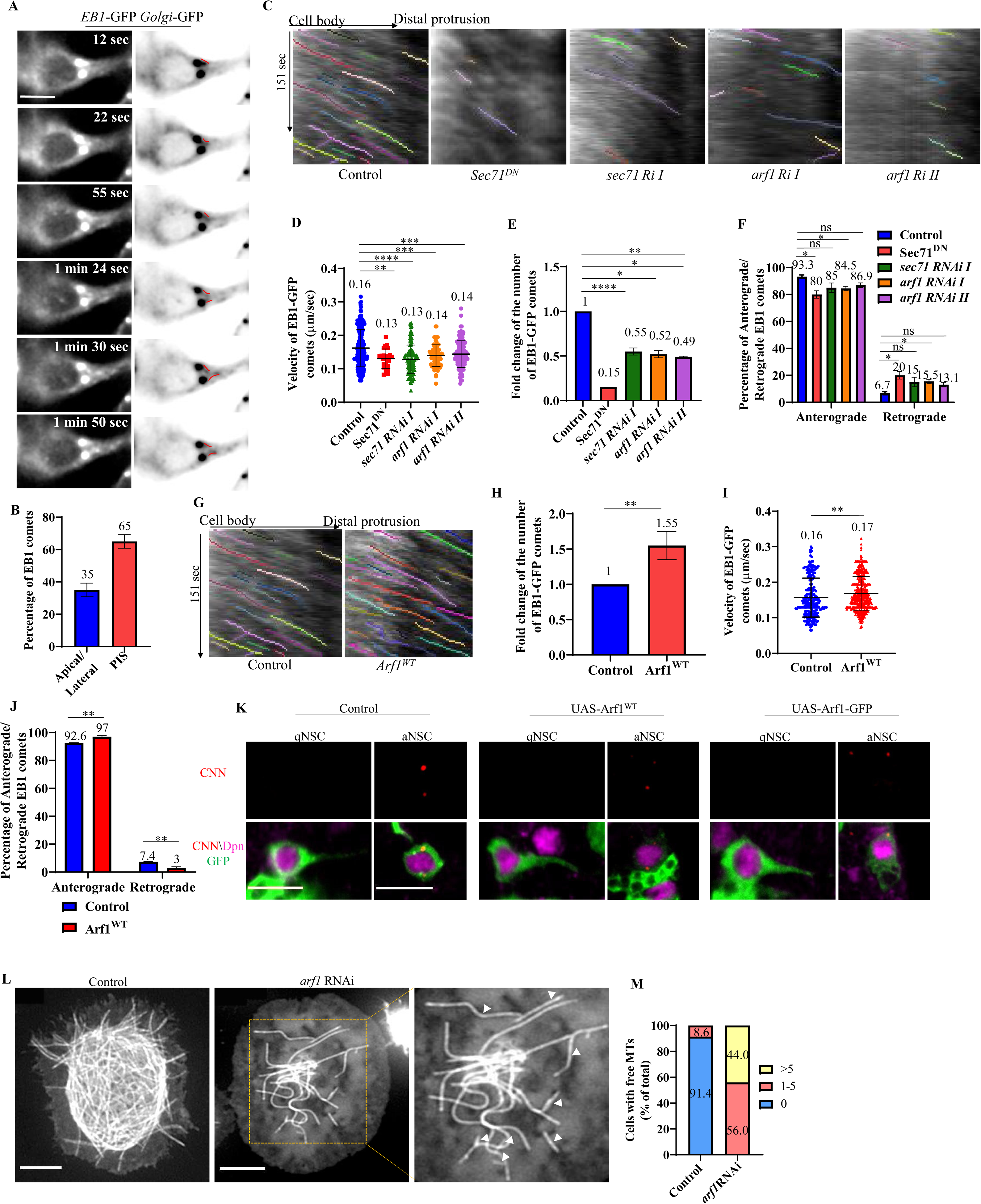
Golgi proteins Arf1 and Sec71 are critical for microtubule growth and orientation in the primary protrusion of qNSCs. A) Time series of a qNSC from larval brains at 6h ALH from *UAS*-*Golgi-GFP* with *grh*-Gal4; *UAS*-EB1-GFP. Red lines indicate the movement of EB1 comets emerging from Golgi present at the PIS region. B) Quantification graph of percentage of EB1 comets arising from the apical/lateral regions vs the PIS region of qNSC from larval brains at 6h ALH from *UAS*-*Golgi-GFP* with *grh*-Gal4; *UAS*-EB1-GFP. EB1 comets emerging from Golgi at the PIS region, 65%, n=228 comets; from Golgi at the apical/lateral region, 35%, n=125 comets, 21 qNSCs. C) Kymographs of EB1-GFP comets movement in the primary protrusion of qNSCs, including the control (*UAS-β-Gal* RNAi), *UAS*-*Sec71^DN^, sec71* RNAi I (*UAS*-*sec71* RNAi /VDRC#100300KK), *arf1* RNAi I (*UAS*-*arf1* RNAi /VDRC#23082GD) and *arf1* RNAi II (*UAS*-*arf1* RNAi/VDRC#103572KK), with EB1-GFP expressed under *grh*-Gal4 at 6h ALH. D) Quantification graph of the velocity of EB1-GFP comets in the primary protrusion of qNSCs at 6h ALH from various genotypes in (C). Control, 0.16, n=270 comets, 32 qNSCs; *Sec71^DN^*, 0.13 fold, n=25 comets, 21 qNSCs; *sec71* RNAi, 0.13, n=126 comets, 24 qNSCs; *arf1* RNAi I, 0.14, n=78 comets, 18 qNSCs; *arf1* RNAi II, 0.14, n=156 comets, 37 qNSCs. ***p<0.001, **p<0.01. E) Quantification graph of fold changes in the number of EB1-GFP comets in the primary protrusion of qNSCs at 6h ALH from various genotypes compared with control in (C). Control, 1, n=270 comets, 32 qNSCs; *Sec71^DN^*, 0.15, n=25 comets, 21 qNSCs; *sec71* RNAi, 0.55, n=126 comets, 24 qNSCs; *arf1* RNAi I, 0.52, n=78 comets, 18 qNSCs; *arf1* RNAi II, 0.49, n=156 comets, 37 qNSCs. ****p<0.0001, **p<0.01, *p<0.05. F) Quantification graph of the percentage of anterograde- and retrograde-moving EB1-GFP comets in the primary protrusion of qNSCs from various genotypes in (C). Anterograde-moving comets: control, 93.3%, n=252 comets; *Sec71^DN^*, 80%, n=20 comets; *sec71* RNAi, 85%, n= 107 comets; *arf1* RNAi I, 84.5%, n=66 comets; *arf1* RNAi II, 85.9%, n=135 comets. Retrograde-moving comets: control, 6.7%, n=18 comets; Sec71^DN^, 20%, n=5 comets; *sec71* RNAi, 15%, n=19 comets; *arf1* RNAi I, 15.5%, n=12 comets; *arf1* RNAi II, 13.1%, n=22 comets. *p<0.05, ns-non-significant. G) Kymograph of EB1-GFP comets movement in the primary protrusion of qNSCs, including the control (*UAS-β-Gal* RNAi) and *UAS*-*Arf1^WT^,* with EB1-GFP expressed under *grh*-Gal4 at 6h ALH. H) Quantification graph of fold changes in the number of EB1-GFP comets in the primary protrusion of qNSCs at 6h ALH from various genotypes compared with control in (G). *Arf1^WT^*, 1.55 fold, n=497 comets, 32 qNSCs, compared with control, 1, n=283 comets, 38 qNSCs. **p<0.01. I) Quantification graph of the velocity of EB1-GFP comets in the primary protrusion of qNSCs at 6h ALH from various genotypes in (G). Control, 0.16 µm/second, n=283 comets, 38 qNSCs; *Arf1^WT^*, 0.17 µm/second, n=497 comets, 32 qNSCs. **p<0.01. J) Quantification graph of the percentage of anterograde- and retrograde-moving EB1-GFP comets in the primary protrusion of qNSCs from various genotypes in (G). Anterograde-moving comets in control, 92.6%, n=262 comets; *Arf1^WT^*, 97%, n=482 comets. Retrograde-moving comets in control, 7.4%, n=21 comets; Arf1^WT^, 3%, n=15 comets. **p<0.01. K) Larval brains at 6h ALH from the control (*UAS-β-Gal* RNAi), *UAS-Arf1^WT^* and *UAS*-*Arf1-GFP* with *grh*-Gal4; *UAS*-CD8-GFP were stained for CNN, Dpn, and GFP. qNSCs show no CNN recruitment (100%, n=34) as compared to activated NSCs (100%, n=21) in all genotypes. L) Time-lapse microscopy of GFP-tubulin wild-type and *arf1*-depleted *Drosophila* S2 cells show that *arf1*-depleted cells have numerous “free” microtubules (white arrow heads) and sparser microtubule network. M) Quantification graph of free microtubules per cell for genotypes in (L). Control, 0 free MTs=91.4%, 1-5 free MTs=8.6%, n=105; *arf1* RNAi, 0 free MTs=0%, 1-5 free MTs=56%, >5 free MTs=44%, n=109. Anterograde movement: from cell body to the tip of the protrusion; Retrograde movement: from the protrusion toward the cell body. Scale bars: 10 μm

### Arf1 and its GEF Sec71 are critical for microtubule assembly and orientation in qNSCs

To further examine the role of the Golgi in acentrosomal microtubule growth, we analyzed the effect of Arf1 and Sec71 on microtubule growth in qNSCs. At 6h ALH, EB1-GFP comets were significantly reduced in the primary protrusion of qNSCs expressing *Sec71^DN^*, compared with robust comets in the control (Figure 4C-F, Movies S9, S10). Furthermore, the velocity of EB1 particles was significantly reduced in *Sec71^DN^* qNSCs, as compared to the control (Figure 4D). These observations suggest that Sec71 is critical for acentrosomal microtubule growth in qNSCs. As we reported previously, most microtubules are oriented “plus-end-out” in qNSCs ^33^; At 6h ALH, 93.3% of EB1-GFP comets in control qNSCs displayed anterograde movement, while the remaining 6.7% displayed retrograde movement (Figure 4F). Interestingly, the percentage of EB1-GFP comets displaying retrograde movement was increased significantly to 20% in *Sec71^DN^* qNSCs (Figure 4F). Similarly, knockdown of *sec71* by RNAi caused a significant reduction in the velocity (0.13 µm/sec in *sec71* RNAi compared with 0.16µm/sec in control) and total number of EB1-GFP comets in qNSCs as compared to the control (Figure 4C-F, Movie S11), as well as a slight but not statistically significant increase in the number of retrograde EB1-GFP comets (Figure 4F). Therefore, Sec71 is critical for acentrosomal microtubule assembly and orientation in qNSCs.

Similarly, *arf1* knockdown by two different RNAi lines showed a significant reduction in velocity and the number of EB1-GFP comets as compared to the control (Figure 4C-F, Movie S9, S12, S13). Loss of *arf1* also caused a significant increase in the number of retrograde EB1-GFP comets (Figure 4F). Conversely, *Arf1* overexpression at 6h ALH resulted in a significantly higher number of EB1-GFP comets in the primary protrusion of qNSCs (Figure 4G-H, Movies S9, and S14). Moreover, the average velocity of EB1-GFP comets was significantly increased upon *Arf1^WT^*overexpression (Figure 4I). In addition, the number of anterograde moving EB1-GFP comets was increased to 97% in Arf1^WT^ as compared to 92.6% in the control (Figure 4J). We wondered whether Arf1 overexpression activates the centrosomes in qNSCs to trigger their premature reactivation. However, this was not the case, as qNSCs at 6h ALH were still devoid of the CDK5RAP2 homologue centrosomin (CNN), an essential component of PCM, upon overexpression of Arf1^WT^ or Arf1-GFP as compared to activated NSCs (Figure 4K). These observations show that Arf1 critically regulates acentrosomal microtubule growth and orientation in qNSCs.

To further elucidate the effect of Arf1 and Sec71 on microtubule orientation, we examined the localization of a microtubule minus-end marker Nod-β-Gal in qNSCs. In control qNSCs, Nod-β-Gal was concentrated at the apical region of the primary protrusion (Figure S4F-G). By contrast, Nod-β-Gal was delocalized from the apical region and distributed around the cell body or PIS region of the primary protrusion upon *arf1* RNAi knockdown, *sec71* RNAi knockdown or *Sec71^DN^* overexpression (Figure S4F-G). Therefore, the microtubule orientation is altered in qNSCs upon *arf1* and *sec71* depletion.

To further demonstrate the importance of Golgi in acentrosomal microtubule growth, we analyzed the effect of Brefeldin A (BFA), a drug disrupts Golgi by inhibiting Arf1 activation ^52,53^, on microtubule growth in qNSCs. At 6h ALH, larval brains treated with BFA showed a dramatic loss of Arf1 or Sec71 puncta as compared to the control brains treated with DMSO (Figure S4H). Remarkably, BFA treatment caused a severe reduction in the velocity and total number of EB1-GFP comets in qNSCs, as well as a significant increase in the number of retrograde EB1-GFP comets (Figure S4I-L, Movies S15, S16).

Taken together, these results indicate that both Arf1 and Sec71 are critical for microtubule growth and plus-end-out orientation in the primary protrusion of qNSCs.

### Arf1 regulates the microtubule network in S2 cells

To further elucidate the importance of Arf1 in microtubule growth and maintenance, we analyzed the effect of loss of *arf1* on the microtubule network by time-lapse imaging of GFP-Tubulin in *Drosophila* S2 cells. In wild-type cells, “free” microtubules (where both the plus and minus ends of the same microtubule are clearly observed) are rarely found ^54^ (Figure 4L-M; Movie S17). In striking contrast, when *arf1* was depleted by RNAi, the microtubule cytoskeleton became less dense and vast majority of cells had free microtubules visible at the cell periphery (Figure 4L-M, Movie S18). Thus, similar to Patronin ^54^, Arf1 regulates the microtubule network in S2 cells.

### Arf1 and Sec71 are required for the regeneration of qNSC protrusion

Our recent work has shown that Msps promotes acentrosomal microtubule growth in qNSCs ^33^. Remarkably, *msps* depletion in NSCs diminished the regeneration ability of qNSC protrusions after injury, even after ∼1 hour of imaging (Figure 5A-B; Movie S19), suggesting that microtubules play an important role in the regeneration process. Since Arf1 and Sec71 regulate microtubule growth in qNSCs (Figure 4), we sought to examine whether these two proteins are required for the regeneration of qNSC protrusions. In *ex vivo* larval brains at 6-8h ALH, only 20% of *Sec71^DN^* qNSCs were able to fully regenerate their protrusions, as compared to 62.5% in control qNSCs, upon laser ablation at the middle region of the protrusion (Figure 5C-D, Movie S2, S20). Similarly, only 11.1% and 33.33% of qNSCs were able to completely regenerate their protrusions upon *arf1* knockdown or *sec71* knockdown following injury, respectively (Figure 5A-D, Movies S21, S22).

**Figure 5.**
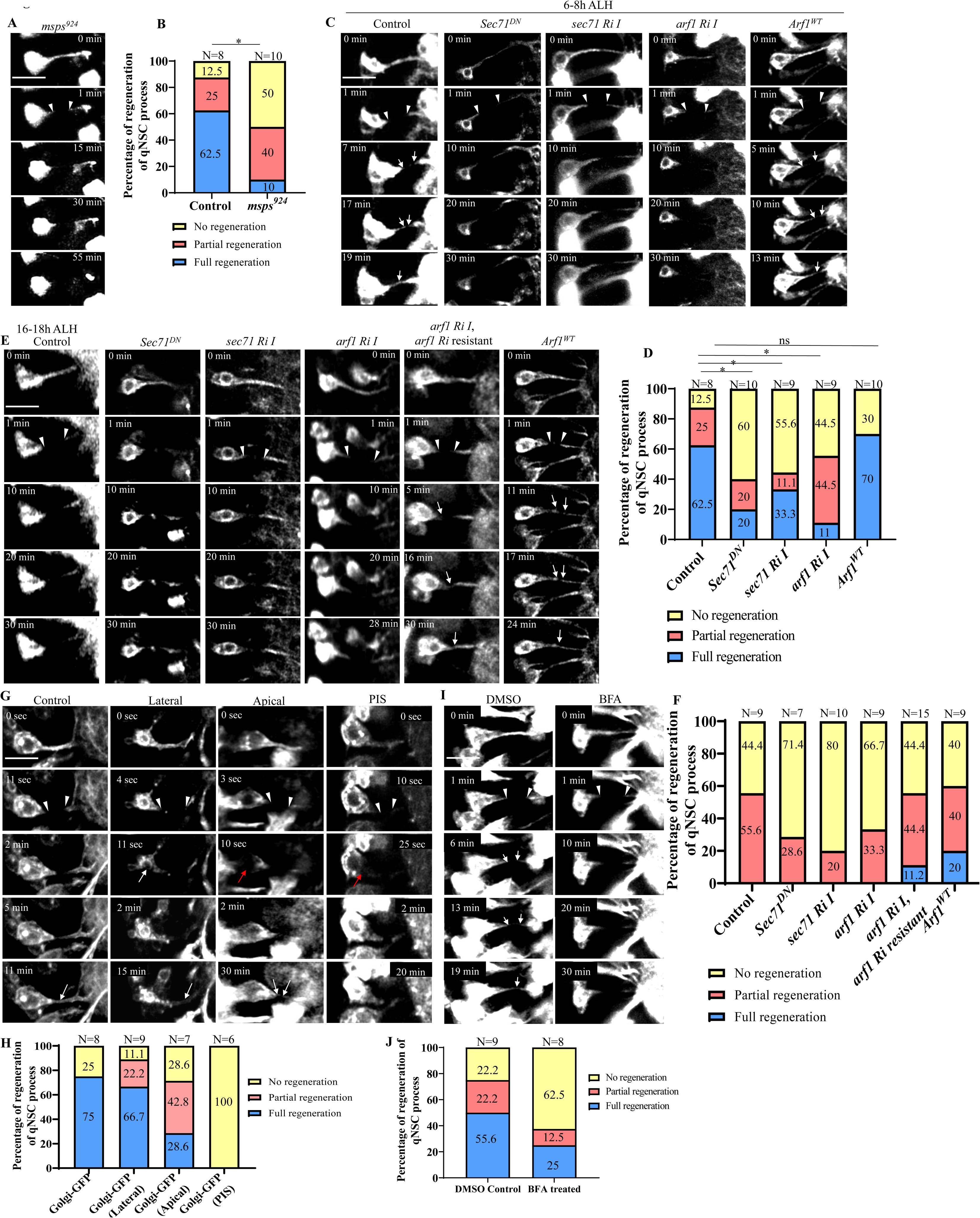
Golgi proteins Arf1 and Sec71 are required for the regeneration of quiescent NSC protrusions upon injury. A) Time series of a qNSC in *ex vivo* larval brain at 6-8h ALH labelled by *grh*-Gal4; *UAS*-mCD8-GFP in *msps^924^*ablated at the middle region of the protrusion. B) Quantification graph of percentage of regeneration of control (*grh*>CD8-GFP; Figure 1C) and *msps^924^* qNSCs expressing *grh*>CD8-GFP at 6-8h ALH after laser ablation. Control, complete regeneration=62.5%, partial regeneration=25%, no regeneration=12.5%, n=8). *msps^924^*, complete regeneration=10%, partial regeneration=40%, no regeneration=50%, n=10. *p< 0.01. C) Time series of a quiescent NSC from *ex vivo* larval brain at 6-8h ALH, including the control, *UAS*-*Sec71^DN^*, *UAS*-*sec71* RNAi I (s*ec71* RNAi (VDRC#100300KK), *UAS*-*arf1* RNAi I (*arf1* RNAi (VDRC#23082GD) and *UAS*-*Arf1^WT^* with *grh*-Gal4; *UAS*-mCD8-GFP ablated at the middle region of the protrusion. D) Quantification graph of the percentage of primary protrusions of qNSCs that are able to regenerate after injury from various genotypes in (C). Control, no regeneration=12.5%, partial regeneration= 25%, complete regeneration=62.5%, n=8; *Sec71^DN^*, no regeneration=60%, partial regeneration= 20%, complete regeneration=20%, n=10; *UAS*-*sec71* RNAi I, no regeneration=55.6%, partial regeneration= 11.1%, complete regeneration=33.3%, n=9; *UAS*-*arf1* RNAi I, no regeneration=44.5%, partial regeneration= 44.5%, complete regeneration=11%, n=9; *Arf1^WT^*, no regeneration=30%, partial regeneration= 0%, complete regeneration=70%, n=10. *p<0.05, ns-non-significant. E) Time series of a quiescent NSC from the larval brain at 16-18h ALH, including the control, *UAS*-*Sec71^DN^*, *UAS*-*sec71* RNAi I (*sec71* RNAi /VDRC#100300KK), *UAS*-*arf1* RNAi I (*arf1* RNAi /VDRC#23082GD), *UAS*-*arf1* RNAi I, *UAS*-*arf1* RNAi I resistant and *UAS*-*Arf1^WT^*, driven by *grh*-Gal4; *UAS*-mCD8-GFP ablated at the middle region of the protrusion. F) Quantification graph of the percentage of primary protrusion of qNSCs that are able to regenerate after injury from various genotypes in (E). Control, no regeneration=44.4%, partial regeneration=55.6% n=9; *Sec71^DN^*, no regeneration =71.4%, partial regeneration= 28.6%, n=7; *UAS*-*sec71* RNAi I, no regeneration=80%, partial regeneration= 20%, n=10; *UAS*-*arf1* RNAi I, no regeneration=66.7%, partial regeneration= 33.3%, n=9; *UAS*-*arf1* RNAi I, *UAS*-*arf1* RNAi I resistant, no regeneration=44.4%, partial regeneration= 44.4%, complete regeneration=11.2%, n=9, *Arf1^WT^*, no regeneration=40%, partial regeneration=40%, complete regeneration=20%, n=15;. *p<0.05, ns- non-significant. G) Time series of regeneration of quiescent NSCs from the larval brain at 6-8h ALH following ablation of Golgi at the lateral, apical, or PIS regions, after the severing of the protrusion at the middle region. Control, ablation at the middle region of the protrusion only. Quiescent NSCs express *UAS*-*mCD8*-GFP and *UAS*-*Golgi*-GFP under the control of *grh*-Gal4. H) Quantification graph of the percentage of qNSCs at 6-8h ALH that can regenerate after laser ablation for genotypes in (G). Control, complete regeneration= 75%, no regeneration=25%, n=8; Lateral Golgi ablation, complete regeneration= 66.7%, no regeneration=11.1%, partial regeneration= 22.2%, n=9; Apical Golgi ablation, complete re-growth= 28.6%, no growth=28.6%, partial growth= 42.8%, n=9; PIS Golgi ablation, no growth=100%, n=6. For ablation of PIS Golgi, only those surviving qNSCs were scored for regeneration. I) Time series of regeneration of quiescent NSCs in larval brains at 6-8h ALH, including control (DMSO-treated) or Brefeldin A (BFA)-treated brains labelled by *grh*-Gal4; *UAS*-mCD8-GFP, ablated at the middle region of the protrusion. J) Quantification graph of the percentage of qNSCs at 6-8h ALH that can regenerate after laser ablation in the control (DMSO-treated) or BFA-treated brains in (I). Control, complete regeneration=55.6%, no regeneration=22.2%, partial growth=22.2%, n=9; BFA-treated, no regeneration=62.5%, partial regeneration=12.5%, Regeneration=25%, n=8. White arrowheads indicate the ablation at the middle region of the protrusion (A, C, E, G, and I); Red arrowheads indicate the ablation at Golgi (G); White arrows indicate regeneration of the qNSC protrusions (A, C, E, G, and I). Scale bars: 10 μm

At 6-8h ALH, Arf1 overexpression did not significantly alter the regeneration capacity in qNSCs (Figure 5C-D, Movies S23, S2 for control). The average time taken for regeneration in both Arf1 overexpressed qNSCs (19.1 minutes) and control qNSCs (19.4 minutes) was also similar. At 16-18h ALH, control qNSCs were unable to regenerate completely, although 55.6% of qNSCs could partially regenerate the severed protrusion (Figure 5E-F, Movies S3). Remarkably, this partial regeneration capacity was diminished in *ex vivo* brains upon *arf1* or *sec71* depletion (Figure 5E-F, Movies S23-26). The regeneration defects observed in *arf1* RNAi I at 16-18h ALH were completely rescued by the co-expression of the *arf1* RNAi-resistant transgene (Figure 5E-F, Movies S27). Strikingly, at 16-18h ALH, Arf1 overexpression in *ex vivo* brains allowed for a complete regeneration in 20% of qNSCs as compared to none in the control (Figure 5E-F, Movies S28, S2 for control). This observation suggests that Arf1 overexpression can enhance the regeneration ability of qNSCs.

Taken together, our data demonstrate that the Golgi proteins Arf1 and its GEF Sec71 are critical for the regeneration of qNSC protrusions, likely through their novel functions in microtubule growth.

### Golgi at the protrusion initial segment is critical for qNSC regeneration

To investigate whether Golgi at the PIS region is important for the regeneration of qNSC protrusions, we performed laser ablation at the Golgi after ablating the middle region of the protrusion. We observed an increased regeneration capacity (75%) in qNSCs co-expressing Golgi-GFP and mCD8-GFP at 6h ALH (Figure 5G-H, Movie S29), compared to control qNSCs expressing mCD8-GFP alone (62.5%, Figure S1C, Movie S2). This might be due to the slight enhancement in microtubule growth caused by Golgi-GFP overexpression (Figure S4D). We then ablated Golgi at the lateral, apical, or PIS regions, following the severing of the protrusion at the middle region. Ablation at lateral region had no significant effect on the regeneration capacity of qNSC protrusions (Figure 5G-H, Movie S30-31). In contrast, disruption of apical Golgi reduced the regeneration capacity of the protrusions, as majority of the protrusions partially regenerated within the 30 min imaging window. (Figure 5G-H, Movie S32-33). Remarkably, disruption of Golgi at the PIS region resulted in a complete failure in the regeneration of qNSC protrusions (Figure 5G-H, Movie S34-35). This data suggests that Golgi at both, the apical, but more importantly at the PIS region are critical for the regeneration capability of qNSC cellular protrusions. Interestingly, ablating the PIS Golgi following the ablation of the cellular protrusion resulted in increased death of qNSCs (45.45%, 5 out of 11), suggesting that the PIS region might also be important for qNSC survival following an injury.

To ascertain the importance of Golgi in the regeneration of qNSC primary protrusions, we performed laser ablation on BFA-treated larval brains at 6h ALH. The localization of Arf1 and Sec71 was completely disrupted after BFA treatment (Figure S4H). When the middle region of primary protrusions was ablated in BFA-treated qNSCs, the protrusions failed to regenerate completely in 75% of qNSCs, as compared to 44.4% of DMSO-treated qNSCs (Figure 5I-J, Movie S36, S37). These results indicate that the Golgi apparatus is required for the regeneration of the primary protrusion of qNSCs after injury.

### Msps physically associates with both GTP- and GDP-bound forms of Arf1

Next, we explored whether Arf1 and Msps can physically interact with each other using proximity ligation assay (PLA), a technique that enables the detection of protein-protein interactions with high specificity and sensitivity ^55^. We co-expressed various proteins tagged with Myc or Flag in S2 cells and quantified PLA foci that indicated protein interactions (Figures 6A-C). The vast majority of S2 cells co-expressing both Flag and Myc controls had no PLA signals, while a fraction of the cells displayed weak PLA fluorescence signal of no more than 10 foci (Figures 6A-C). Similarly, the vast majority of cells (82.7-89%) co-expressing Myc-Msps with control Flag or Flag-Arf1^WT^ with control Myc had no PLA signal; 0.9 and 0.71 PLA foci per cell, were detected under each co-expression condition, respectively (Figures 6A-C). By contrast, 90% of cells co-expressing Flag-Arf1^WT^ and Myc-Msps displayed PLA signal (15.4 PLA foci per cell on average), of which 19.7% displayed strong signal (>30 foci), 35% displayed moderate signal (11-30 foci), and 35.3% displayed weak signal (1-10 foci) (Figures 6B-C). Further, we tested whether Msps associates with the GTP- or GDP-bound form of Arf1. In controls expressing either Myc and HA, Myc-Msps and HA, Myc and HA-Arf1^Q71L^ (Arf1-GTP) or Myc and HA-Arf1^T31N^ (Arf1-GDP), we observed that the majority of cells had no PLA signal (Figures S5A-C). However, the majority of S2 cells co-expressing either Myc-Msps and HA-Arf1^Q71L^ or Myc-Msps and HA-Arf1^T31N^ showed good PLA signals (Figures S5A-C), suggesting that Msps physically associates with both GTP- and GDP-bound forms of Arf1.

**Figure 6.**
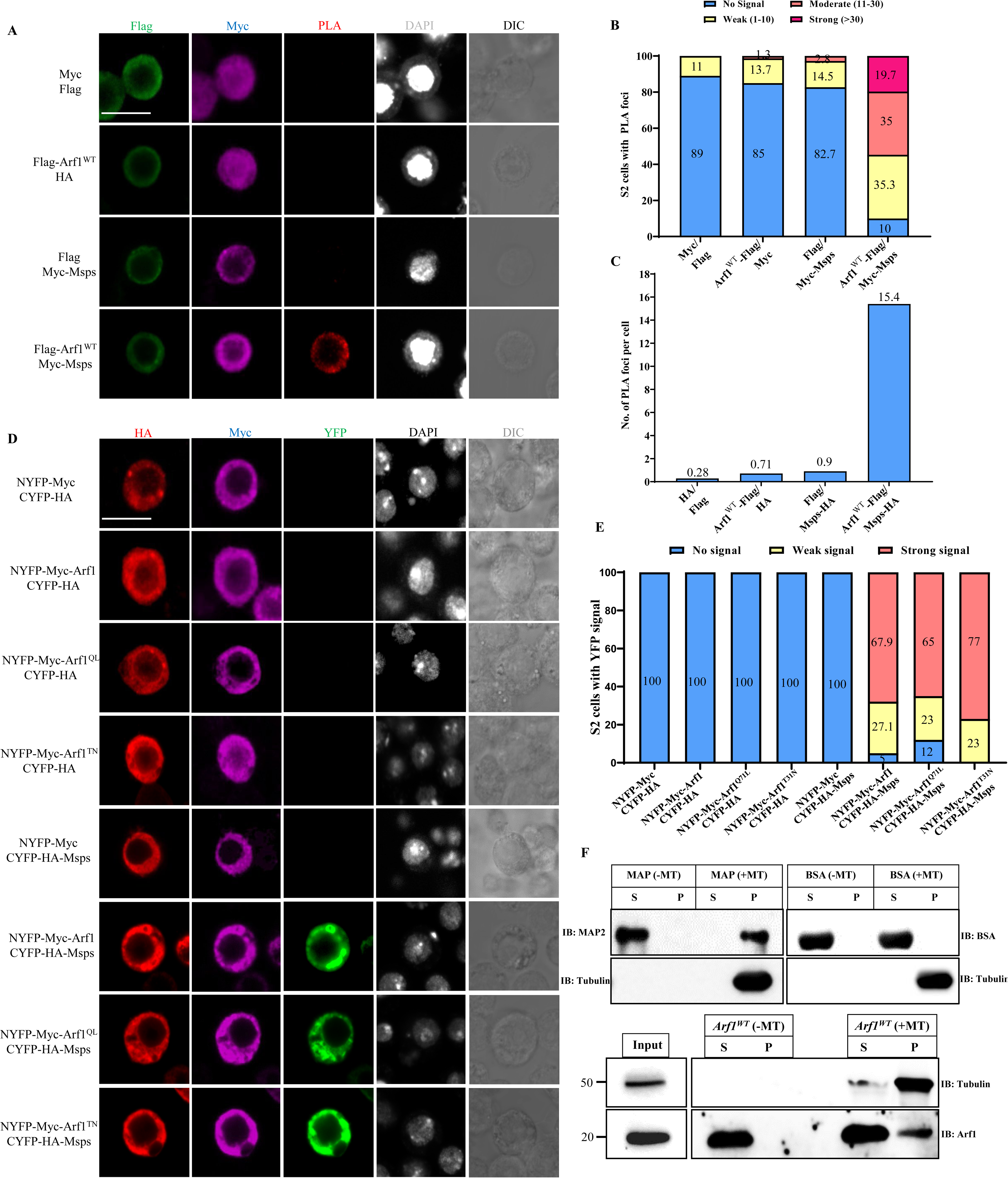
The Golgi protein Arf1 physically associates with Msps. (A) *In situ* PLA assay between Arf1^WT^ and Msps in S2 cells. S2 cells transfected with two of the indicated plasmids (Myc, Flag, Flag-Arf1^WT^, Myc-Msps,) were stained for Flag, Myc, and DNA and detected for PLA signal. Cell outlines were determined by differential interference contrast (DIC) images. (B) Quantification graphs showing the percentage of S2 cells with no PLA signal, weak (≤10 foci), moderate (11–30 foci), and strong (>30 foci) PLA signals for (A). Myc and Flag, no signal=89%, weak signal=11%, n=215; Flag-Arf1^WT^ and Myc, no signal=85%, weak=13.7%, moderate=1.3%, n=222; Flag and Myc-Msps, no signal=82.7%, weak=14.5%, moderate=2.8%, n=271; Flag-Arf1^WT^ and Myc-Msps, no signal=10%, weak=35.3%, moderate=35%, strong=19.7%, n=247. (C) Quantification graph of the average number of PLA foci per cell in (A). Myc and Flag, 0.28, n=215; Flag-Arf1^WT^ and Myc, 0.71, n=222; Flag and Myc-Msps, 0.9, n=271; Flag-Arf1^WT^ and Myc-Msps, 15.4, n=247. (D) *In vitro* BiFC assay between Msps and Arf1 (WT, Q71L and T31N). S2 cells that were triple transfected with *actin*-Gal4, *UAS*-CYFP-HA-Msps (or *UAS*-CYFP-HA as a control), and *UAS*-NYFP-Myc-Arf1 (or *UAS*-NYFP-Myc-Arf1^Q71L^ or *UAS*-NYFP-Myc-Arf1^T31N^ or *UAS*-NYFP-Myc as a control) were stained with Myc (blue) and HA (red) and detected for YFP fluorescence (green). Cell outlines were observed using differential interference contrast (DIC) imaging. (E) Quantification graph of the percentage of S2 cells with no YFP signal, weak YFP signal and strong YFP signal for (D). NYFP-Myc and CYFP-HA, no YFP signal=100%, n=122; NYFP-Myc-Arf1^WT^ and CYFP-HA, no YFP signal=100%, n=78; NYFP-Myc-Arf1^Q71L^ and CYFP-HA, no YFP signal=100%, n=143; NYFP-Myc-Arf1^T31N^ and CYFP-HA, no YFP signal=100%, n=126; NYFP-Myc and CYFP-HA-Msps, no YFP signal=100%, n=107; NYFP-Myc-Arf1^WT^ and CYFP-HA-Msps, no YFP signal=5%, weak YFP signal=27.1%, strong YFP signal=67.9%, n=81; NYFP-Myc-Arf1^Q71L^ and CYFP-HA-Msps, no YFP signal=12%, weak YFP signal=23%, strong YFP signal=65%, n=70 and NYFP-Myc-Arf1^T31N^ and CYFP-HA-Msps; no YFP signal=0%, weak YFP signal=23%, strong YFP signal=77%, n=100. (F) Microtubules were polymerized *in vitro* and added to protein extracts from whole embryos overexpressing Arf1 under *grh*-Gal4. Binding reactions with (+MT) or without (−MT) microtubules were stratified on a 20% sucrose cushion. The resulting pellets (P) and supernatants (S) were analyzed by Western analysis. Embryo extracts incubated with or without MTs were probed with antibodies specific for Arf1 and α-tubulin. Arf1 cosedimented with microtubules. The microtubule associated protein (MAP2) and bovine serum albumin (BSA) were used as positive and negative controls respectively. Scale bars: 5 μm

To validate the association between Msps and Arf1, we employed another protein-protein interaction assay named biomolecular fluorescence complementation (BiFC), which can detect transient or weak interactions due to the irreversibility of the BiFC complex formation ^56,57^. We generated the following chimeric proteins: CYFP-HA-Msps (Msps with C-terminal YFP tagged with HA), NYFP-Myc-Arf1^WT^, NYFP-Myc-Arf1^Q71L^, and NYFP-Myc-Arf1^T31N^ (N-terminal YFP tagged with Myc was fused to Arf1^WT^, Arf1^Q71L^ or Arf1^T31N^). As expected, no YFP signal was detected in S2 cells that were transfected with either of these two chimeric constructs and their respective controls, NYFP-Myc and CYFP-HA (Figure 6D-E). By contrast, strong YFP signal was detected when cells were transfected with both NYFP-Myc-Arf1^WT^ and CYFP-HA-Msps (Figure 6D-E). Similarly, strong YFP signal was detected in cells co-expressing either NYFP-Myc-Arf1^Q71L^ and CYFP-HA-Msps or NYFP-Myc-Arf1^T31N^ and CYFP-HA-Msps, supporting that Msps physically associates with both GTP- and GDP-bound forms of Arf1. Therefore, Msps is likely an effector of Arf1 during NSC reactivation.

### Arf1 associates with microtubules *in vitro*

Since Arf1 affected microtubule growth *in vivo* and *in vitro* (Figure 4) and can physically associate with the microtubule polymerase Msps (Figure 6A-E), we next examined whether Arf1 can cosediment with microtubules as a complex *in vitro*^58,59^. The microtubule associated protein (MAP2) was used as a positive control wherein it cosedimented with microtubules in the pellet fraction (Figure 6F). Bovine serum albumin (BSA) which does not associate with microtubules was used as a negative control (Figure 6F). Supernatant from whole embryos overexpressing Arf1 under the control of *grh*-Gal4 driver were incubated with *in vitro* polymerized microtubules. A significant fraction of Arf1 cosedimented with microtubules in the pellet fraction in the presence of taxol-stabilized bovine microtubules (Figure 6F). In the absence of microtubules, Arf1 is found in the supernatant but not in the pellet (Figure 6F). This data suggests that Arf1 is able to associate with microtubules *in vitro*.

### Arf1-Msps-E-cad pathway promotes NSC reactivation

Recently, we showed that the cell-adhesion molecule E-cadherin/Shotgun localizes to the NSC-neuropil contact sites in an Msps-dependent manner and is required for NSC reactivation (Deng et al., 2021). To determine the role of functional Golgi in this process, we examined E-cad localization and intensity at the NSC-neuropil contact sites in *arf1* and *sec71* loss-of-function conditions. Strikingly, at 16h ALH, basal NSC-neuropil contact sites were lost in 53.3% of *arf1* RNAi I and 60% of *arf1* RNAi II qNSCs, compared with their loss in 12.7% of control qNSCs (Figure 7A, C). Likewise, *sec71* knockdown using two different RNAi lines led to E-cad delocalization from NSC-neuropil contact sites in 60.6% and 44.7% of qNSCs, respectively, compared with E-cad delocalization in 12.7% of control cells (Figure 7B-C). The intensity of E- cad at NSC-neuropil contact sites (basal E-Cad) in the ventral nerve cord (VNC) was dramatically reduced in *arf1* and *sec71* knockdown as compared to control qNSCs (Figure 7A-B, D). The depletion of either *arf1* or *sec71* led to a significant reduction in the ratio of basal E-cad intensity to E-cad intensity at the cellular process, supporting the loss of enrichment of E-cad at the NSC-neuropil contact sites (Figure 7A, B, E). Interestingly, although Arf1 overexpression did not alter the percentage of E-cad basal localization at NSC-neuropil contact sites, it resulted in a small but significant increase in the E-cad levels at the NSC-neuropil contact sites (Figure 7F-H). Therefore, E-cad localization to NSC-neuropil contact sites requires functional Golgi proteins Arf1 and Sec71 within NSCs.

**Figure 7.**
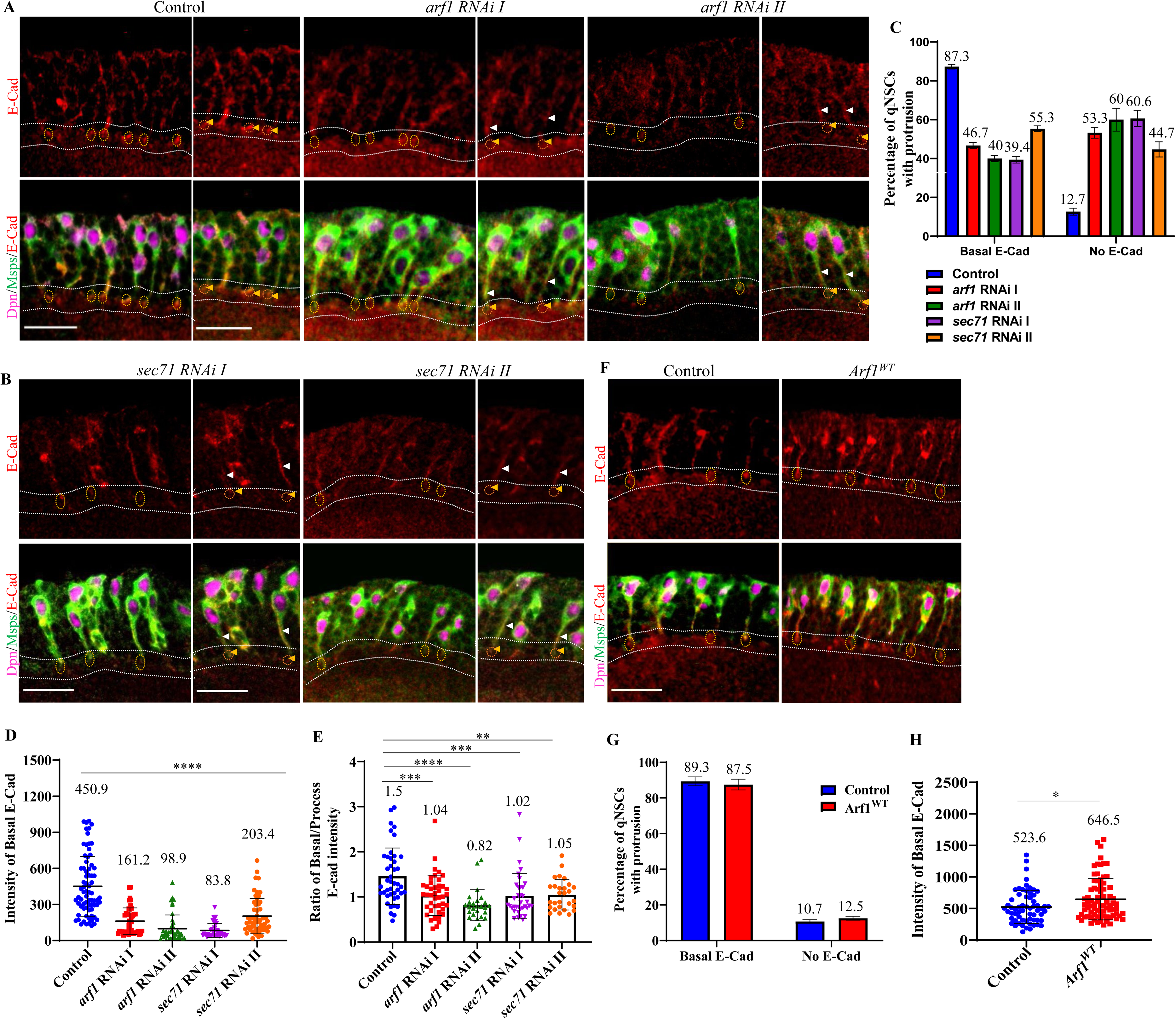
Arf1 and Sec71 target E-cad to the NSC-neuropil contact sites. A) The larval ventral nerve cord (VNC) at 16h ALH, including the control (*UAS-β-Gal* RNAi), *UAS*-*arf1* RNAi I (VDRC#23082GD), and *UAS*-*arf1* RNAi II (VDRC#103572KK) driven by *grh*-Gal4, were labelled with E-cad, Dpn, and Msps. The primary protrusions of qNSCs were marked by Msps. B) The larval ventral nerve cord (VNC) at 16h ALH, including the *UAS*-*sec71* RNAi I (VDRC#100300KK) and *UAS*-*sec71* RNAi II (BSDC#32366) driven by *grh*-Gal4, were labelled with E-cad, Dpn, and Msps. The primary protrusions of qNSCs were marked by Msps. C) Quantification of E-Cadherin basal localization at putative NSC-neuropil contact sites in qNSCs from genotypes in (A and B). “No E-cad” means absent or strongly reduced E-cad, whereas “Basal E-cad” denotes localization at the NSC-neuropil contact sites. Basal E-cad localization: control, 83.3%, n=118 NSCs; *arf1* RNAi I, 46.7%, n=107 NSCs; *arf1* RNAi II, 40%, n=95 NSCs; *sec71* RNAi I, 39.4%, n=99 NSCs and *sec71* RNAi II, 55.3%, n=67 NSCs. ***p<0.001. D) Quantification graph of basal E-Cadherin intensity at putative NSC-neuropil contact sites in qNSCs from genotypes in (A and B). Control, 450.9 A.U., n=73 NSCs; *arf1* RNAi I, 161.2 A.U., n=46 NSCs; *arf1* RNAi II, 98.9 A.U., n=44 NSCs; *sec71* RNAi I, 83.8 A.U., n=44 NSCs; *sec71* RNAi II, 203.4 A.U., n=54 NSCs. ****p<0.0001. E) Quantification graph of the ratio of basal E-Cadherin intensity at NSC-neuropil contact sites to E-cadherin intensity along the protrusion in qNSCs from genotypes in (A and B). Control, 1.5, n=42 NSCs; *arf1* RNAi I, 1.04, n=41 NSCs; *arf1* RNAi II, 0.82, n=26 NSCs; *sec71* RNAi I, 1.02, n=36 NSCs and *sec71* RNAi II, 1.05, n=28 NSCs. **p<0.01, ***p<0.001, ****p<0.0001. F) The larval ventral nerve cord (VNC) at 16h ALH, including the control (*UAS-β-Gal* RNAi) and *UAS*-*Arf1^WT^* driven by *grh*-Gal4, were labelled with E-cad, Dpn, and Msps. The primary protrusions of qNSCs were marked by Msps. G) Quantification of E-Cadherin basal localization at putative NSC-neuropil contact sites in qNSCs from genotypes in (F). “No E-cad” means absent or strongly reduced E-cad, whereas “Basal E-cad” denotes localization at the NSC-neuropil contact sites. Basal E-cad localization: control, 89.3%, n=94 NSCs; *Arf1^WT^*, 87.5%, n=88 NSCs. H) Quantification graph of basal E-Cadherin intensity at NSC-neuropil contact sites in qNSCs from genotypes in (F). Control, 523.6 A.U., n=62 NSCs; *Arf1^WT^*, 646.5 A.U., n=78 NSCs. *p<0.05.

Next, we sought to investigate the epistasis of *arf1*, *msps* and *E-cad* during NSC reactivation. First, we overexpressed *msps* in *arf1* and *sec71-*depleted brains and found a strong suppression of NSC reactivation phenotypes. At 24 h ALH, the number of EdU-negative NSCs in *arf1*–depleted brains overexpressing *Msps^FL^* was dramatically reduced to 17% compared with 46.5% in *arf1* RNAi control brains (Figure 8A-B). Remarkably, overexpression of *msps* in *arf1*-depleted qNSC also significantly suppressed the loss of EB-1 comet number, velocity, and orientation defects caused by *arf1* RNAi (Figure 8C-F, Movies S38-41). In contrast, over-expression of *Arf1* could not suppress the NSC delayed reactivation phenotype observed in in *msps*-depleted brains (Figure S6A-B). Furthermore, in S2 cells, Msps overexpression significantly suppressed the loss of microtubule density and number of free microtubules that were caused by *arf1* RNAi knockdown (Figure 8G-H, Movies S42-44). These genetic data indicate that Msps is a major effector of Arf1 during NSC reactivation and microtubule growth.

**Figure 8.**
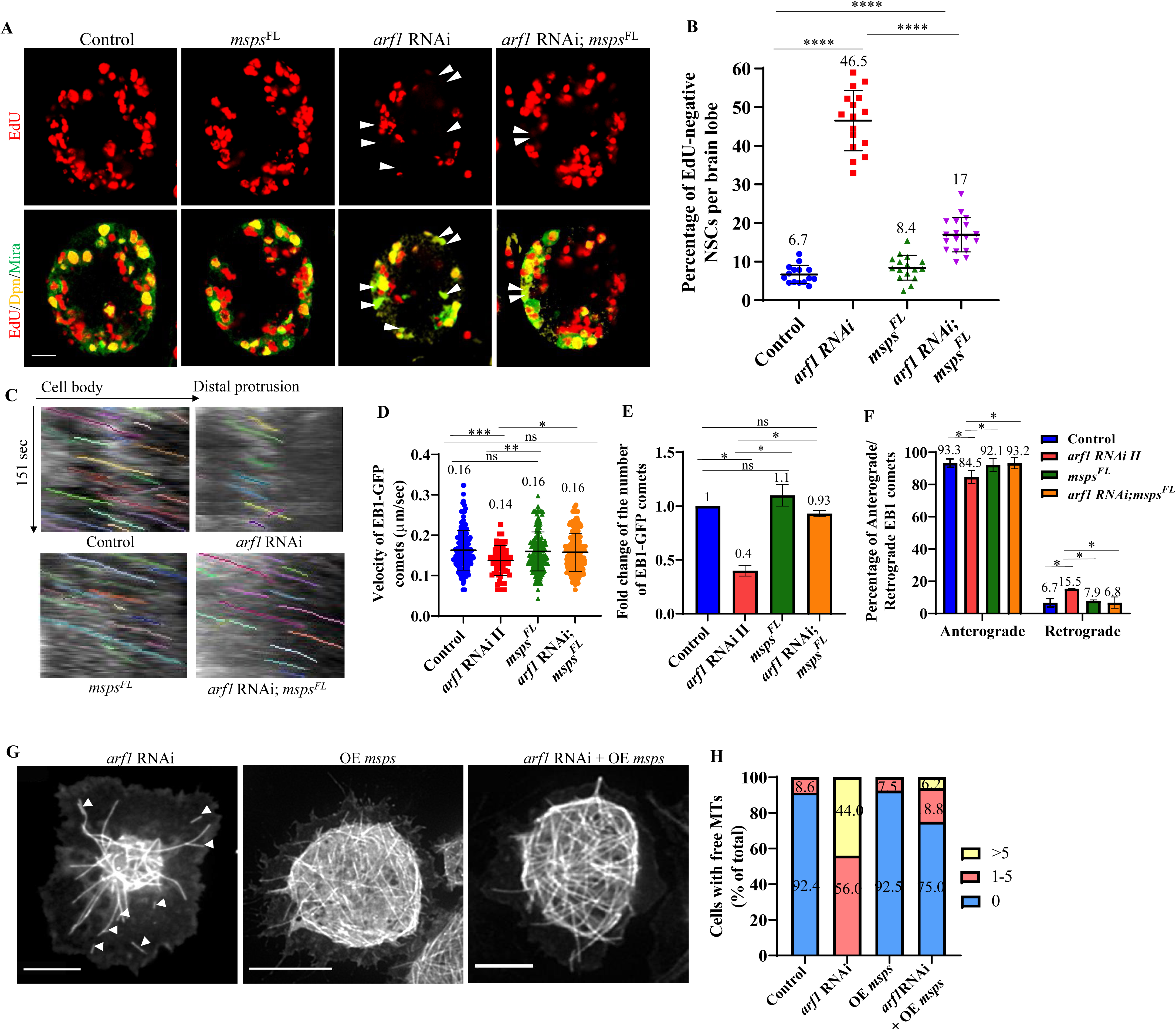
The microtubule regulator Msps and cell adhesion molecule E-cad act downstream of Arf1 to promote NSC reactivation. A) Larval brains of various genotypes at 24h ALH were analysed for EdU incorporation. Control: *grh*-Gal4; *UAS*-*Dcr2*/*UAS-β-Gal* RNAi. *msps^FL^*: *grh*-Gal4 *β-Gal* RNAi, *UAS*-*msps^FL^*. *arf1* RNAi: *grh*-Gal4 *β-Gal* RNAi, *UAS*-*arf1* RNAi II (VDRC#103572KK). *arf1* RNAi; *msps^FL^*: *grh*-Gal4 *UAS*-*arf1* RNAi II; *UAS*-*msps^FL^,* NSCs were marked by Dpn and Mira. White arrowheads point to NSCs without EdU incorporation. B) Quantification graph of EdU-negative NSCs per brain lobe for genotypes in (A). Control, 6.7%, n=15 BL; *arf1* RNAi, 46.5%, n=16; *msps^FL^*, 8.4%, n=16 BL; and *arf1* RNAi; *msps^FL^*, 17%, n=18 BL. ****p<0.0001. C) Kymograph of EB1-GFP comets movement in the primary protrusion of qNSCs of various genotypes in (A). EB1-GFP was expressed under *grh*-Gal4 at 6h ALH. D) Quantification graph of the velocity of EB1-GFP comets in the primary protrusion of qNSCs at 6h ALH from various genotypes in (C). Control, 0.16, n=165 comets, 18 qNSCs; *arf1* RNAi II, 0.14, n=65 comets, 20 qNSCs; *msps^FL^*, 0.16 fold, n=204 comets, 18 qNSCs and *arf1* RNAi II; *msps^FL^*, 0.16, n=250 comets, 24 qNSCs. ***p<0.001, **p<0.01, *p<0.05 ns- non-significant. E) Quantification graph of fold changes in the number of EB1-GFP comets in the primary protrusion of qNSCs at 6h ALH from various genotypes compared with control in (C). Control, 1, n=165 comets, 18 qNSCs; *arf1* RNAi II, n=65 comets, 20 qNSCs; *msps^FL^*, 1.1, n= n=204 comets, 18 qNSCs and *arf1* RNAi II; *msps^FL^*, 0.93, n=250 comets, 24 qNSCs. **p<0.01, *p<0.05. F) Quantification graph of the percentage of anterograde- and retrograde-moving EB1-GFP comets in the primary protrusion of qNSCs from various genotypes in (C). Anterograde-moving comets: control, 93.3%, n=154 comets; *arf1* RNAi II, 84.5%, n=55 comets; *msps^FL^*, 92.1%, n= 188 comets; *arf1* RNAi II; *msps^FL^*, 93.2%, n=233 comets. Retrograde-moving comets: control, 6.7%, n=11 comets; *arf1* RNAi II, 15.5%, n=10 comets, *msps^FL^*, 7.9%, n= 16 comets; *arf1* RNAi II; *msps^FL^*, 6.8%, n=17 comets. **p<0.01, *p<0.05, ns- non-significant. G) Time-lapse microscopy of GFP-tubulin in *arf1*-depleted, over-expression *msps*, and *arf1*-depleted + over-expression *msps Drosophila* S2 cells. H) Quantification graph of free microtubules per cell for genotypes in (G). Control, 0 free MTs=91.4%, 1-5 free MTs=8.6%, n=105; *arf1* RNAi, 0 free MTs=0%, 1-5 free MTs=56%, >5 free MTs=44%, n=109; OE *msps*, 0 free MTs=92.5%, 1-5 free MTs=7.5%, n=93 and *arf1* RNAi + OE *msps*, 0 free MTs=75%, 1-5 free MTs=18.8%, >5 free MTs=6.2%, n=64. Scale bars: 10 μm

Similarly, in *sec71*–depleted brains overexpressing *Msps^FL^*, the number of EdU-negative NSCs was significantly reduced to 13.2% compared with 36.7% in *sec71* RNAi brains (Figure S6C-D). Next, we overexpressed E-cad in NSCs with *arf1* depletion and tested its ability to suppress NSC reactivation defects. At 24 h ALH, significantly fewer EdU-negative NSCs were observed in *arf1* RNAi knockdown with *E-cad* overexpressed brains than in *arf1*-depleted brains (Figure S6E-F). Therefore, E-cad overexpression suppressed NSC reactivation defects caused by *arf1* RNAi knockdown. Further, *E-cad; arf1* double knockdown resulted in significantly more NSCs with failed EdU incorporation, compared to single knockdown or RNAi control (Figure S6G-H). Taken together, these data strongly support a role for the Arf1-Msps-E-cad genetic pathway in promoting NSC reactivation.

## Discussion

In this study, we have discovered that *Drosophila* quiescent NSC protrusions can regenerate upon injury. We have identified two critical Golgi proteins—a small GTPase Arf1 and its GEF Sec71/Arf1GEF—as key regulators of acentrosomal microtubule assembly, qNSC reactivation, and regeneration upon injury. We have also shown that Golgi in qNSCs acts as the acentrosomal microtubule organizing center of qNSCs. We propose that Arf1 physically associates with Msps, a microtubule polymerase and a new effector of Arf1 during acentrosomal microtubule growth and qNSC reactivation. Finally, Arf1 functions upstream of Msps to target the cell-adhesion molecule E-cad to NSC-neuropil contact sites during qNSC reactivation. Taken together, our data has established *Drosophila* qNSCs as a new regeneration model and identified a novel Arf1-Msps pathway in the regulation of microtubule growth and qNSC reactivation.

### Regeneration of qNSCs: a new regeneration model

While it is well established that most axons of the adult mammalian central nervous system (CNS) fail to regenerate after an injury, it is unknown whether NSCs have the ability to regrow their fibres. This work demonstrates that *Drosophila* qNSCs can repair and regenerate their primary protrusions upon injury. This new regeneration model shares at least two common features with other well-known regeneration models, such as adult rat retinal ganglion cells (RGCs) and adult *C. elegans* motor neurons: first, *Drosophila* qNSCs exhibit age-dependent decline in regeneration capability. We found that the regeneration capability is the greatest in *Drosophila* qNSCs derived from newly hatched larvae and declines within the 24h duration of quiescence during development. This is similar to age-dependent decline in axon regeneration capacity after axotomy in *C. elegans* mechanosensory neurons and in corticospinal and rubrospinal neurons in the adult mammalian CNS ^60,61^. Second, alterations in microtubule dynamics affect regeneration. We demonstrated that the microtubule regulators Arf1, Sec71, and Msps are required for the regeneration of *Drosophila* qNSCs upon injury, highlighting the significance of microtubule growth in the regeneration process. In addition, the level of stable microtubules in qNSCs, marked by acetylated-tubulin, decreased over time during development, which correlates with the decline in the regeneration capability of these cells. The effects of microtubule dynamics on qNSC regeneration is in line with findings in the mammalian CNS neurons that microtubule manipulation promotes axon regeneration after injury ^62–64^. One interesting aspect of *Drosophila* qNSCs regeneration is that the distal tip of the protrusion does not degenerate after severing, but is capable of the regeneration similar to the proximal end. Likely, certain cues or signals from the neuropil plays an important role in this regeneration process, as the distal tip of the protrusion is attached to the neuropil. Our findings on *Drosophila* qNSC regeneration highlight qNSCs as a valuable model to identify novel factors as well as the underlying mechanisms that promote CNS regeneration.

### Golgi functions as the acentrosomal microtubule organizing centre in qNSCs

The microtubules in the primary protrusion of qNSCs are predominantly acentrosomal; the qNSCs in newly hatched larvae possess inactive centrosomes, which mature over time during NSC reactivation ^33^. Although Golgi has emerged as potential acentrosomal MTOCs in several cell types ^43,52,65–67^, the role of Golgi in acentrosomal microtubule growth in qNSC was unknown. We have revealed that Golgi are present as 1-3 punctate structures at the PIS and pericentrosomal regions, as compared to the scattered distribution of smaller Golgi puncta in the cytosol and perinuclear region of dividing NSCs. The majority of microtubule nucleation sites, marked by the emerging EB1-GFP comets, were found to be at the Golgi in the PIS region (from where primary protrusions are extended from the cell body). Interestingly, microtubule nucleation sites are also found at apical Golgi where some EB1-GFP comets move towards the PIS Golgi. Microtubules originating at the apical Golgi might be severed to generate shorter microtubules that function as seeds for nucleation and growth at the PIS Golgi. As Golgi are often pericentrosomal in qNSCs and the centrosome is immature in these qNSCs, the centrosomes may take over Golgi as the MTOC when NSCs re-enter the cell cycle. Although the exact mechanism is unknown, the proximity between the immature centrosomes and the apical Golgi likely facilitates this transition.

Furthermore, we have reported that the apical region and more importantly the PIS region containing Golgi is required for the regeneration of qNSC protrusions upon injury, likely via microtubule growth. The disruption of Golgi functions by *arf1*/*sec71* depletion or Brefeldin A treatment also abolished the regeneration capability of qNSCs. In addition, the critical role played by the microtubule polymerase Msps in qNSC regeneration also supports the notion that microtubule growth is required for qNSC regeneration upon injury. Similarly, alterations in microtubule dynamics is important for axon regeneration. Moderately stable microtubules are required for efficient axon regeneration, as treatment with low doses of microtubule-stabilizing agent taxol or epothilone B enhanced axon regeneration in central neurons following spinal cord injury ^63,68,69^. Such moderate stabilization of microtubules helps to shift microtubule polymerization towards the axon tip and enables growth cone formation in adult injured CNS neurons, thereby improving their regeneration ^63,69^. The anterograde transport of cargo, such as mitochondria, vesicles, and proteins, along the microtubules is also important for their delivery to lesioned axonal tip for axon regeneration ^70,71^. Further studies are needed to better understand how microtubule growth and dynamics enhances qNSC regeneration.

### Arf1 plays a novel role in regulating microtubule growth via its effector Msps/XMAP215

The Golgi-resident small GTPase Arf1 facilitates vesicle formation and trafficking at Golgi, processes that involve the actin cytoskeleton and actin-binding proteins ^72–76^. Although Golgi-derived vesicles are transported along microtubules in various cell types such as fibroblasts and neurons ^77^, the role of Arf1 in regulating microtubule functions was unknown. In this study, we uncovered a novel role for Arf1 and its GEF Sec71/Arf1GEF in regulating microtubule growth. Specifically, we have identified Arf1 and Sec71 as key regulators of acentrosomal microtubule nucleation, growth, and orientation in the primary protrusion of *Drosophila* qNSCs. *arf1* or *sec71* loss-of-function resulted in reduced microtubule growth and misorientation, while Arf1 overexpression enhanced microtubule growth and the anterograde movement of EB1 comets in qNSCs. Besides qNSCs, Arf1 also promotes microtubule growth and stabilization of microtubule network in *Drosophila* S2 cells, as shown by *in vitro* time-lapse imaging. The novel role of Arf1 in regulating microtubule growth is unexpected, as it is well-established that Arf1-6 proteins are critical for membrane trafficking, while Arf-like (Arl) proteins such as Arl2 are important for microtubule functions. Interestingly, mammalian Arf1 recruits tubulin-binding cofactor E (TBCE), which is known to associate with the microtubule regulator Arl2 during microtubule polymerization, to the Golgi and Arf1 overexpression increases tubulin abundance in motor neurons ^78^. It will be particularly interesting to investigate a similar microtubule regulatory role for Arf1 in other *Drosophila* cell types as well as in different organisms.

Furthermore, the following findings have led us to infer that Msps is an effector of Arf1. First, depletion of *arf1*, *sec71* or *msps* led to similar phenotypes in microtubule growth and orientation and in NSC reactivation. Second, Arf1, Sec71 and Msps were all required for qNSC regeneration upon injury. Third, both GTP- and GDP-bound forms of Arf1 physically associated with Msps. Arf1-GDP, which is present in the cytosol, is converted to the GTP-bound form by its GEF such as Sec71 upon recruitment to the Golgi membrane ^46^. Therefore, Msps seems to be associated with Arf1 both in the cytosol and at the Golgi. Finally, Msps overexpression largely rescued defects in NSC reactivation and microtubule growth and orientation caused by *arf1*-depletion. It is interesting to note that this new role of Arf1 in NSC reactivation appears to be independent of its well-known function in membrane trafficking, as COPI and AP-1 are not important for NSC reactivation. By identifying Msps as a new effector of Arf1, our study further validates an important role for Arf1 in regulating microtubule growth. Overall, we have identified the Arf1-Msps-E-cad axis as a key pathway that promotes qNSC cell cycle re-entry during reactivation.

Taken together, our study has established *Drosophila* qNSCs as a new regeneration model and has identified a critical role for the Golgi proteins Arf1 and Sec71 in promoting acentrosomal microtubule growth, reactivation, and regeneration of qNSCs. Furthermore, it demonstrates how Arf1 and its new effector Msps-mediated acentrosomal microtubule growth facilitates the targeting of E-cad to NSC-neuropil contact sites to promote NSC reactivation. Our findings may serve as a general paradigm that could be applied to other types of quiescent stem cells in both *Drosophila* and mammalian systems.

## Materials and Methods

### Fly stocks and genetics

Fly stocks and genetic crosses were raised at 25°C unless otherwise stated. Fly stocks were kept in vials or bottles containing standard fly food (0.8% Drosophila agar, 5.8% Cornmeal, 5.1% Dextrose, and 2.4% Brewer’s yeast). The following fly strains were used in this study: *insc*-Gal4 (BDSC#8751; 1407-Gal4), *grh*-Gal4 (A. Brand), *insc*-Gal4 tub-Gal80ts, *msps*^924^ (F. Yu), *UAS*t-*Arf1^Q71L^* (F. Yu), *UAS*t-*ArfI^WT^* (F. Yu), *UAS*t-*ArfI^DN^* (F. Yu), *UAS*t-*Sec71^DN^* (F. Yu), *UAS*t-*Sec71^WT^*(F. Yu), *UAS*t-*Arf1-*GFP (F. Yu), *UAS*t-*ManII*-Venus (F. Yu), *UAS*p-*Nod-β-gal* (F. Yu), *UAS*t-*arf1* RNAi resistant (F. Yu), *UAS*t-*sec71* RNAi resistant (F. Yu), *UAS*t-*Msps^FL^* and *UAS*p-*EBI*-GFP (F. Yu).

The following stocks were obtained from Bloomington Drosophila Stock Center (BDSC): *UAS*-*sec71* RNAi (BDSC#32366), *UAS*-*E-cad* RNAi (BDSC#32904), *UAS*-*E-cad* RNAi (BDSC#38207), *UAS*-*AP-1-2β* RNAi (BDSC#31707), *UAS*-*AP-1-2β* RNAi (BDSC#28328), *UAS*-*AP-1γ* RNAi (BDSC#23533), *UAS*-*AP-1µ* RNAi (BDSC#27534), *UAS*-*AP-1σ* RNAi (BDSC#40895), *UAS*-*αCOP1* RNAi (BDSC#31702), *UAS*-*βCOP1* RNAi (BDSC#33741), *UAS*-*δCOP1* RNAi (BDSC#31764), *UAS*-*εCOP1* RNAi (BDSC#28890), *UAS*-*γCOP1* RNAi (BDSC#28889), *UAS*-*ΖCOP1* RNAi (BDSC#17808) and *UAS*-Golgi-GFP (BDSC#32902). The following stocks were obtained from Vienna Drosophila Resource Center (VDRC): *UAS*-*arf1* RNAi (23082GD), *UAS*-*arf1* RNAi (103572KK), *UAS*-*sec71* RNAi (100300KK) and *UAS*-*msps* RNAi (21982GD). *UAS*-*β-Gal* RNAi (BDSC#50680) is often used as a control *UAS* element to balance the total number of *UAS* elements in each genotype. The knockdown efficiency of all *arf1* and *sec71* RNAi lines in larval brains has been verified by immunostaining with anti-Arf1 and anti-Sec71 antibodies. The various RNAi knockdown or overexpression constructs were induced using *grh*-Gal4 or *insc*-Gal4 unless otherwise stated.

All experiments were carried out at 25°C, except for RNAi knockdown or overexpression studies that were performed at 29°C, unless otherwise indicated.

### EdU (5-ethynyl-2’-deoxyuridine) incorporation assay

Larvae of various genotypes were fed with food supplemented with 0.2 mM EdU from ClickiT® EdU Imaging Kits (Invitrogen) for 4 h. The larval brains were dissected in PBS and fixed with 4% EM-grade formaldehyde in PBS for 22 min, followed by washing thrice (each wash for 10 min) with 0.3% PBST, and blocked with 3% BSA in PBST for 30 min. The incorporated EdU was detected by Alexa Fluor azide, according to the Click-iT EdU protocol (Invitrogen). The brains were rinsed twice and subjected to standard immunohistochemistry.

### Immunohistochemistry

*Drosophila* larvae were dissected in PBS, and the larval brains were fixed in 4% EM-grade formaldehyde in PBT (PBS + 0.3% Triton-100) for 22 min. The samples were processed for immunostaining as previously described (Li et al., 2017). For α-tubulin immunohistochemistry, the larvae were dissected in Shield and Sang M3 medium (Sigma-Aldrich) supplemented with 10% FBS, followed by fixation in 10% formaldehyde in Testis buffer (183 mM KCl, 47 mM NaCl, 10 mM Tris-HCl, and 1 mM EDTA, pH 6.8) supplemented with 0.01% Triton X-100). The fixed brains were washed once in PBS and twice in 0.1% Triton X-100 in PBS. Images were taken using LSM710 confocal microscope system (Axio Observer Z1; ZEISS), fitted with a PlanApochromat 40×/1.3 NA oil differential interference contrast objective, and brightness and contrast were adjusted by Photoshop CS6.

The primary antibodies used in this paper were guinea pig anti-Dpn (1:1,000), rabbit anti-Dpn (1:200), mouse anti-Mira (1:50, F. Matsuzaki), rabbit anti-GFP (1:3,000; F. Yu), mouse anti-GFP (1:3000; F. Yu), rabbit anti-RFP (1:2000; abcam, Cat#62341), guinea pig anti-Asl (1:200, C. Gonzalez), rabbit anti-Sas-4 (1:100, J. Raff), mouse anti-α-tubulin (1:200, Sigma, Cat#: T6199), mouse anti-γ-tubulin (1:200, Sigma, Cat#: T5326), rabbit anti-CNN (1:5,000, E. Schejter and T. Megraw), rabbit anti-Msps (1:500), rabbit anti-Msps (1:1,000, J. Raff), rabbit anti-PH3 (1:200, Sigma, Cat#: 06-570), rat anti-E-cadherin (1:20, DCAD2, DSHB), mouse anti-β-galactosidase (1:1,000, Promega, Cat#: Z3781), rabbit anti-β-galactosidase (1:5,000, Invitrogen, A-11132), mouse anti-Sec71 (1:100, F. Yu), rabbit anti-GM130 (1:200, Abcam ab52649), guinea pig anti-Arf1 (1:200, F. Yu), rabbit anti-D-TACC (1:200), rabbit anti-Insc (1:1,000), rabbit anti-Flag (1:1,000; Sigma-Aldrich), mouse anti-Myc (1:1,000; abcam), rabbit anti-HA (1:1000, Sigma-Aldrich) and Topro (1:5000). The secondary antibodies used were conjugated with Alexa Fluor 488, 555 or 647 (Jackson laboratory).

### Laser ablation of qNSCs

Larval brains of various genotypes expressing *UAS*-mCD8-GFP under *grh*-Gal4 at various time points were dissected in Shield and Sang M3 insect medium (Sigma-Aldrich) supplemented with 10% FBS. The *ex vivo* larval brain explant culture was supplied with fat body from wild-type third instar and live imaging of the larval brains were performed with a Nikon A1R MP laser scanning confocal microscope using 40X objective lens and Zoom factor 5. Quiescent NSCs with protrusions attached to the neuropil were chosen and imaged for about 10-30 seconds (1sec/frame) before ablation. qNSCs were hit by a picolaser emitting 70-80nW of laser power for 1-1.5 secs to cause injury ^79^. After injury, qNSCs were imaged again for 10-30 seconds followed by time-lapse imaging for 30 minutes (1min/frame, 7-10 z-stacks with 0.5-0.8-µm intervals). The movies and images were made and analysed with NIH ImageJ software.

*In vivo* imaging and ablation were done using whole larvae, which were placed in a single layer PDMS microfluidic device, vacuum was applied via a syringe to immobilize the animal (Mishra et al., 2014).

### Tracking of EB1-GFP comets

Larval brains of various genotypes expressing EB1-GFP under *grh*-Gal4 at various time points were dissected in Shield and Sang M3 insect medium (Sigma-Aldrich) supplemented with 10% FBS. The larval brain explant culture was supplied with fat body from wild-type third instar and live imaging of the larval brains were performed with LSM710 confocal microscope system using 40X Oil lens and Zoom factor 6. The brains were imaged for 151 seconds, with 83 frames acquired for each movie and the images were analysed with NIH ImageJ software. The velocity of the EB1-GFP comets were calculated and kymographs were generated using KymoButler ^80^. For EB1 comets arising from Golgi, the larval brains were imaged for 90 seconds with 50 frames acquired for each movie. The tracking of EB1 comets were done manually using NIH ImageJ software.

### Brefeldin A (BFA)-treatment of *Drosophila* larval brains

Larval brains expressing *UAS*-mCD8-GFP under *grh*-Gal4 at 6h ALH were dissected in Shield and Sang M3 insect medium (Sigma-Aldrich) supplemented with 10% FBS. The brains were then incubated in either 10 μg/ml BFA (Sigma-Aldrich) dissolved in DMSO or the same volume of DMSO only as negative control in Shield and Sang M3 insect medium (Sigma-Aldrich) supplemented with 10% FBS. Following BFA treatment for 30 minutes, live imaging of EB1-GFP tracking or laser ablation experiments were performed. For Arf1 and Sec71 antibody detection after the treatments, the larval samples were fixed and immunostained as described above.

### Microtubule Pulldown Assay

Determination of the ability of Arf1 to cosediment with exogenously added microtubules was performed according to the manufacturer’s protocol (Cytoskeleton, Denver) with some modifications ^58^. Protein was extracted from whole embryos using an extraction buffer containing 0.1 M Pipes, pH 6.9, 0.9 M glycerol, 5 mM EGTA, 2.5 mM MgSO4 in the presence of a cocktail of protease’s inhibitors (Roche Diagnostics). The extract was clarified by centrifugation at 5000 × g for 20 min at 4°C. The protein content of the supernatant was determined according to the Bradford assay and equivalent amounts of proteins were incubated with *in vitro* polymerized microtubules or without microtubules, following the protocol proposed by the manufacturer. Both reactions were stratified on a 20% sucrose cushion and centrifuged at 37,000 rpm (rotor TL100) for 45 min. The resulting supernatants and pellets were analyzed by Western analysis.

### Cell lines and transfection

*Drosophila* S2 cells (CVCL_Z232) originally from William Chia’s laboratory (with a non-authenticated identity but have been used in the laboratory for the past 10 years) were cultured in Express Five serum-free medium (Gibco) supplemented with 2 mM Glutamine (Thermo Fisher Scientific). The S2 cell culture used in this study is free of mycoplasma contamination, inferred by the absence of small speckles of DAPI staining outside of the cell nucleus. For transient expression of proteins, S2 cells were transfected using Effectene Transfection Reagent (QIAGEN) according to the manufacturer’s protocol. S2 cells were used for RNAi, PLA and BiFC assays.

For RNAi experiments, *Drosophila* S2 cells were cultured and incubated with dsRNA and GFP-α-tubulin plasmid (Gohta Goshima) as previously described ^54,81^. Here, cells were treated with dsRNA for 2 days and analyzed at day 5.

### Bimolecular fluorescence complementation

In vitro bimolecular fluorescence complementation (BiFC) assay (Gohl et al., 2010) was performed using S2 cells. 1 × 10^6^ cells were seeded onto Poly-L-lysine-coated coverslips (Iwaki), and were transfected with *act*-Gal4 and the BiFC constructs each at 0.2 μg per well, respectively, using Effectene Transfection Reagent (QIAGEN). The following BiFC constructs were used to test interactions between Msps and Arf1: *UAS*t-NYFP-Myc-Arf1, *UAS*t-NYFP-Myc-Arf1^Q71L^, *UAS*t-NYFP-Myc-Arf1^T31N^, *UAS*t-CYFP-HA, *UAS*t-NYFP-Myc and *UAS*t-CYFP-HA-Msps. *UAS*t-NYFP-Myc-Arf1, *UAS*t-NYFP-Myc-Arf1^Q71L^, *UAS*t-NYFP-Myc-Arf1^T31N^ and *UAS*t-CYFP-HA-Msps, to test interaction between Msps and Arf1. At 48 hrs after transfection, the growth medium was removed and the cells were rinsed with cold PBS before fixing them with 4% EM grade formaldehyde in PBS for 15 min. The fixed S2 cells were rinsed three times with PBS-T (1xPBS + 0.1% Triton-X100) and blocked with 5% BSA in PBS-T for 1 hr, before incubating them with primary antibodies at room temperature (RT) for 2 hr. Following incubation, the cells were rinsed three times with PBS-T and were incubated with secondary antibodies in PBS-T for 1 hr at RT. Coverslips coated with immuno-stained S2 cells were mounted on to glass slides using vector shield (Vector Laboratory) for confocal microscopy.

### Proximity ligation assay (PLA)

PLA is based on the following principle: Secondary antibodies conjugated with a PLA PLUS or PLA MINUS probe bind to anti-Flag and anti-Myc antibodies, respectively. During ligation, connector oligos hybridize to PLA probes and T4 ligases catalyze to form a circularized template. DNA polymerase amplifies the circularized template, which is bound by fluorescently-labeled complementary oligos, allowing the interaction to be observed as PLA foci within the cells (Adopted from Duolink PLA, Merck). PLA was performed on S2 cells that were transfected with the following plasmids using Effectene Transfection Reagent (QIAGEN): control Myc, control Flag, control HA, Flag-Arf1^WT^, HA-Arf1^Q71L^, HA-Arf1^T31N^ and Myc-Msps. The cells were washed thrice with cold PBS, fixed with 4% EM-grade formaldehyde in PBS for 15 min and blocked in 5% BSA in PBS-T (0.1% Triton-X100) for 45min. The cells were then incubated with primary antibodies at RT for 2 hrs before proceeding with Duolink PLA (Sigma-Aldrich) according to the manufacturer’s protocol. After incubation with primary antibodies, the cells were incubated with PLA probes at 37°C for 1 hr. They were then washed twice with Buffer A for 5 min, each at RT, followed by ligation of probes at 37°C for 30 min. Amplification was performed at 37°C for 100 min, followed by two washes with Buffer B, each for 10 min, at RT. The cells were washed once with 0.01x Buffer B before incubating with primary antibodies diluted in 3% BSA in PBS for 2 hrs at RT. Following this, the cells were washed twice with 0.1% PBS-T and incubated with secondary antibodies for 1.5 hrs at RT, before mounting them with in situ mounting media with DAPI (Duolink, Sigma-Aldrich).

### Quantification and statistical analysis

*Drosophila* larval brains from various genotypes were placed dorsal side up on confocal slides. Confocal z-stacks were taken from the surface to the deep layers of the larval brains (20-30 slides per z-stack with 2 or 3 µm intervals). For each genotype, at least 5 brain lobes were imaged for z-stacks and Image J or Zen softwares were used for quantification.

Statistical analysis was performed using GraphPad Prism 8. Unpaired two-tailed t-tests were used for the comparison of two sample groups and one-way ANOVA or two-way ANOVA followed by Sidak’s multiple comparisons test for the comparison of more than two sample groups. All data are shown as mean ± SD. Statistically non-significant (ns) denotes P > 0.05, * denotes P < 0.0001. All experiments were performed with a minimum of two repeats. In general, n refers to number of NSCs counted, unless otherwise indicated.

## Supporting information

All supplementary movies

## Acknowledgements

We thank F. Yu, Gohta Goshima, Marc Freeman, C. Gonzalez, J. Raff, T. Lee, F Matsuzaki, W. Chia, and the Bloomington Drosophila Stock Center, Vienna Drosophila Resource Center, Kyoto Stock Centre DGGR, and the Developmental Studies Hybridoma Bank for fly stocks and antibodies. We thank Ravinuthula Sruthi Jagannathan for her critical comments on this manuscript. This work is supported by the Ministry of Health-Singapore National Medical Research Council MOH-000143 (MOH-OFIRG18may-0004) to H.W. and Ministry of Education Tier 2 MOE-T2EP30220-0016 to Y.T.

## Author contributions

Conceptualization, MG and HW; Methodology, Data curation, and formal analysis, MG, YG, XT, QD, YST; Writing-original draft, MG and HW; Writing-review & editing, HW, MG, YT, YG, QD; funding acquisition, HW, YT; Resources, HW, YT; Supervision, HW.

## Supplementary Movie Legends

**Movie S1.** Time-lapse imaging of regeneration of the primary protrusion of a qNSC expressing *grh*-Gal4; *UAS*-mCD8-GFP in *ex vivo* larval brain at 0-2h ALH after laser ablation from. Time-lapse before and during ablation is 1 sec/frame and after ablation is 1 min/frame. Time scale: minute: second and hour: minute. Scale bar: 10 µm.

**Movie S2.** Time-lapse imaging of regeneration of the primary protrusion of a qNSC expressing *grh*-Gal4; *UAS*-mCD8-GFP in larval brain *in vivo* at 0-2h ALH after laser ablation. Time scale: minute: second and hour: minute. Scale bar: 10 µm.

**Movie S3.** Time-lapse imaging of cell death after laser ablation at the cell body of a qNSC in *in vivo* larval brain expressing *grh*-Gal4; *UAS*-mCD8-GFP at 6-8h ALH. Time scale: minute: second and hour: minute. Scale bar: 10 µm.

**Movie S4.** Time-lapse imaging of regeneration of the primary protrusion of a qNSC expressing *grh*-Gal4; *UAS*-mCD8-GFP in *ex vivo* larval brain at 6-8h ALH after laser ablation. Time scale: minute: second and hour: minute. Scale bar: 10 µm.

**Movie S5.** Time-lapse imaging of failed regeneration of the primary protrusion of a qNSC expressing *grh*-Gal4; *UAS*-mCD8-GFP in *ex vivo* larval brain at 16-18h ALH after laser ablation. Time scale: minute: second and hour: minute. Scale bar: 10 µm.

**Movie S6.** Time-lapse imaging of failed regeneration of the primary protrusion of a qNSC expressing *grh*-Gal4; *UAS*-mCD8-GFP in *ex vivo* larval brain at 6-8h ALH after laser ablation at PIS region. Time scale: minute: second and hour: minute. Scale bar: 10 µm.

**Movie S7.** Time-lapse imaging of EB1-GFP comets arising from Golgi at PIS region in a qNSC in the larval brain at 6h ALH. *UAS*-*Golgi-GFP* and *UAS*-EB1-GFP were driven by *grh*-Gal4. Time scale: minute: second. Scale bar: 10 µm.

**Movie S8.** Time-lapse imaging of EB1-GFP comets arising from Golgi at the apical or lateral regions in a qNSC in the larval brain at 6h ALH. *UAS*-*Golgi-GFP* and *UAS*-EB1-GFP were driven by *grh*-Gal4. Time scale: minute: second. Scale bar: 10 µm.

**Movie S9.** Time-lapse imaging of EB1-GFP comets in the primary protrusion of a qNSC in the larval brain expressing *grh*-Gal4; *UAS*-EB1-GFP *with UAS*-*β-Gal* RNAi at 6h ALH. Time scale: minute: second. Scale bar: 10 µm.

**Movie S10.** Time-lapse imaging of EB1-GFP comets in the primary protrusion of a qNSC in the larval brain expressing *UAS*-*Sec71^DN^* and *UAS*-EB1-GFP driven by *grh*-Gal4 at 6h ALH. Time scale: minute: second. Scale bar: 10 µm.

**Movie S11.** Time-lapse imaging of EB1-GFP comets in the primary protrusion of a qNSC in the larval brain expressing *UAS*-*sec71* RNAi I (VDRC#100300KK) with *grh*-Gal4; *UAS*-EB1-GFP at 6h ALH. Time scale: minute: second. Scale bar: 10 µm.

**Movie S12.** Time-lapse imaging of EB1-GFP comets in the primary protrusion of a qNSC in the larval brain expressing *UAS*-*arf1* RNAi I (VDRC#23082GD) with *grh*-Gal4; *UAS*-EB1-GFP at 6h ALH. Time scale: minute: second. Scale bar: 10 µm.

**Movie S13.** Time-lapse imaging of EB1-GFP comets in the primary protrusion of a qNSC in the larval brain expressing *UAS*-*arf1* RNAi II (VDRC#103572KK) with *grh*-Gal4; *UAS*-EB1-GFP at 6h ALH. Time scale: minute: second. Scale bar: 10 µm.

**Movie S14.** Time-lapse imaging of EB1-GFP comets in the primary protrusion of a qNSC in the larval brain expressing *UAS*-*Arf1^WT^* with *grh*-Gal4; *UAS*-EB1-GFP at 6h ALH. Time scale: minute: second. Scale bar: 10 µm.

**Movie S15.** Time-lapse imaging of EB1-GFP comets in the primary protrusion of a qNSC in the larval brain expressing *grh*-Gal4; *UAS*-EB1-GFP treated with DMSO at 6h ALH. Time scale: minute: second. Scale bar: 10 µm.

**Movie S16.** Time-lapse imaging of EB1-GFP comets in the primary protrusion of a qNSC in the larval brain expressing *grh*-Gal4; *UAS*-EB1-GFP treated with BFA at 6h ALH. Time scale: minute: second. Scale bar: 10 µm.

**Movie S17.** Time-lapse imaging of GFP-Tubulin in control S2 cells. Time scale: minute: second. Scale bar: 10 µm.

**Movie S18.** Time-lapse imaging of GFP-Tubulin in S2 cells treated with dsRNA *arf1*. Time scale: minute: second. Scale bar: 10 µm.

**Movie S19.** Time-lapse imaging of failed regeneration of the primary protrusion of a qNSC in an *msps^924^* larval brain with *grh*-Gal4; *UAS*-mCD8-GFP at 6-8h ALH after laser ablation. Time scale: minute: second and hour: minute. Scale bar: 10 µm.

**Movie S20.** Time-lapse imaging of failed regeneration of the primary protrusion of a qNSC in the larval brain expressing *UAS*-*Sec71^DN^*with *grh*-Gal4; *UAS*-mCD8-GFP at 6-8h ALH after laser ablation. Time scale: minute: second and hour: minute. Scale bar: 10 µm.

**Movie S21.** Time-lapse imaging of failed regeneration of the primary protrusion of a qNSC in the larval brain expressing *UAS*-*sec71* RNAi I (VDRC#100300KK) with *grh*-Gal4; *UAS*-mCD8-GFP at 6-8h ALH after laser ablation. Time scale: minute: second and hour: minute. Scale bar: 10 µm.

**Movie S22.** Time-lapse imaging of failed regeneration of the primary protrusion of a qNSC in the larval brain expressing *UAS*-*arf1* RNAi I (VDRC#23082GD) with *grh*-Gal4; *UAS*-mCD8-GFP at 6-8h ALH after laser ablation. Time scale: minute: second and hour: minute. Scale bar: 10 µm.

**Movie S23.** Time-lapse imaging of regeneration of the primary protrusion of a qNSC in the larval brain expressing *UAS*-*Arf1^WT^*with *grh*-Gal4; *UAS*-mCD8-GFP after laser ablation at 6-8h ALH. Time scale: minute: second and hour: minute. Scale bar: 10 µm.

**Movie S24.** Time-lapse imaging of failed regeneration of the primary protrusion of a qNSC in the larval brain expressing *UAS*-*Sec71^DN^*with *grh*-Gal4; *UAS*-mCD8-GFP at 16-18h ALH after laser ablation. Time scale: minute: second and hour: minute. Scale bar: 10 µm.

**Movie S25.** Time-lapse imaging of failed regeneration of the primary protrusion of a qNSC in the larval brain expressing *UAS*-*sec71* RNAi I (*sec71* RNAi (VDRC#100300KK) with *grh*-Gal4; *UAS*-mCD8-GFP at 16-18h ALH after laser ablation. Time scale: minute: second and hour: minute. Scale bar: 10 µm.

**Movie S26.** Time-lapse imaging of failed regeneration of the primary protrusion of a qNSC in the larval brain expressing *UAS*-*arf1* RNAi I (VDRC#23082GD) with *grh*-Gal4; *UAS*-mCD8-GFP at 16-18h ALH after laser ablation. Time scale: minute: second and hour: minute. Scale bar: 10 µm.

**Movie S27.** Time-lapse imaging of regeneration of the primary protrusion of a qNSC in the larval brain expressing *UAS*-*arf1* RNAi I, *UAS*-*arf1* RNAi I resistant with *grh*-Gal4; *UAS*-mCD8-GFP at 16-18h ALH after laser ablation. Time scale: minute: second and hour: minute. Scale bar: 10 µm.

**Movie S28.** Time-lapse imaging of regeneration of the primary protrusion of a qNSC in the larval brain expressing *UAS*-*Arf1^WT^* with *grh*-Gal4; *UAS*-mCD8-GFP at 16-18h ALH after laser ablation. Time scale: minute: second and hour: minute. Scale bar: 10 µm.

**Movie S29.** Time-lapse imaging of laser ablation at the middle region of the primary protrusion of a qNSC in the larval brain expressing *UAS*-*Golgi-GFP* with *grh*-Gal4; *UAS*-mCD8-GFP at 6-8h ALH. Time scale: minute: second. Scale bar: 10 µm.

**Movie S30.** Time-lapse imaging of laser ablation at lateral region of the cell body following protrusion severing at the middle region of the primary protrusion of a qNSC in the larval brain expressing *UAS*-*Golgi-GFP* with *grh*-Gal4; *UAS*-mCD8-GFP at 6-8h ALH. Time scale: minute: second. Scale bar: 10 µm.

**Movie S31.** Time-lapse imaging of full regeneration after laser ablation at lateral region of the cell body following protrusion severing at the middle region of the primary protrusion of a qNSC in the larval brain expressing *UAS*-*Golgi-GFP* and *UAS*-mCD8-GFP by *grh*-Gal4 at 6-8h ALH. Time scale: hour: minute. Scale bar: 10 µm.

**Movie S32.** Time-lapse imaging of laser ablation at the apical Golgi following protrusion severing at the middle region of the primary protrusion of a qNSC in the larval brain expressing *UAS*-*Golgi-GFP* with *grh*-Gal4; *UAS*-mCD8-GFP at 6-8h ALH. Time scale: minute: second. Scale bar: 10 µm.

**Movie S33.** Time-lapse imaging of partial regeneration after laser ablation in the primary protrusion of qNSCs in larval brains at 6-8h ALH from *UAS*-*Golgi-GFP* with *grh*-Gal4; *UAS*-mCD8-GFP. Time scale: hour: minute. Scale bar: 10 µm.

**Movie S34.** Time-lapse imaging of laser ablation at the PIS Golgi following protrusion severing at the middle region of the primary protrusion of a qNSC in the larval brain expressing *UAS*-*Golgi-GFP* with *grh*-Gal4; *UAS*-mCD8-GFP at 6-8h ALH. Time scale: minute: second. Scale bar: 10 µm.

**Movie S35.** Time-lapse imaging of failed regeneration after laser ablation at the PIS Golgi following protrusion severing at the middle region of the primary protrusion of a qNSC in the larval brain expressing *UAS*-*Golgi-GFP* with *grh*-Gal4; *UAS*-mCD8-GFP at 6-8h ALH. Time scale: hour: minute. Scale bar: 10 µm.

**Movie S36.** Time-lapse imaging of regeneration after laser ablation in the primary protrusion of a qNSC in the larval brain expressing *grh*-Gal4; *UAS*-mCD8-GFP treated with DMSO. Time scale: minute: second and hour: minute. Scale bar: 10 µm.

**Movie S37.** Time-lapse imaging of failed regeneration after laser ablation in the primary protrusion of a qNSC in the larval brain expressing *grh*-Gal4; *UAS*-mCD8-GFP treated with BFA. Time scale: minute: second and hour: minute. Scale bar: 10 µm.

**Movie S38.** Time-lapse imaging of EB1-GFP comets in the primary protrusion of a qNSC in the larval brain from *UAS*-*β-Gal* RNAi; *UAS*-*β-Gal* with *grh*-Gal4; *UAS*-EB1-GFP at 6h ALH. Time scale: minute: second. Scale bar: 10 µm.

**Movie S39.** Time-lapse imaging of EB1-GFP comets in the primary protrusion of a qNSC in the larval brain from *UAS*-*arf1* RNAi II (VDRC#103572KK); *UAS*-*β-Gal* with *grh*-Gal4; *UAS*-EB1-GFP at 6h ALH. Time scale: minute: second. Scale bar: 10 µm.

**Movie S40.** Time-lapse imaging of EB1-GFP comets in the primary protrusion of a qNSC in the larval brain from *UAS*-*β-Gal; UAS*-*Msps^FL^* with *grh*-Gal4; *UAS*-EB1-GFP at 6h ALH. Time scale: minute: second. Scale bar: 10 µm.

**Movie S41.** Time-lapse imaging of EB1-GFP comets in the primary protrusion of a qNSC in the larval brain from *UAS*-*arf1* RNAi II (VDRC#103572KK);*UAS*-*Msps^FL^* with *grh*-Gal4; *UAS*-EB1-GFP at 6h ALH. Time scale: minute: second. Scale bar: 10 µm.

**Movie S42.** Time-lapse imaging of GFP-Tubulin in S2 cells treated with dsRNA *arf1*. Time scale: minute: second. Scale bar: 10 µm.

**Movie S43.** Time-lapse imaging of GFP-Tubulin in S2 cells treated with OE *msps*. Time scale: minute: second. Scale bar: 10 µm.

**Movie S44.** Time-lapse imaging of GFP-Tubulin in S2 cells treated with dsRNA *arf1* and OE *msps*. Time scale: minute: second. Scale bar: 10 µm.

---

**Figure S1.**
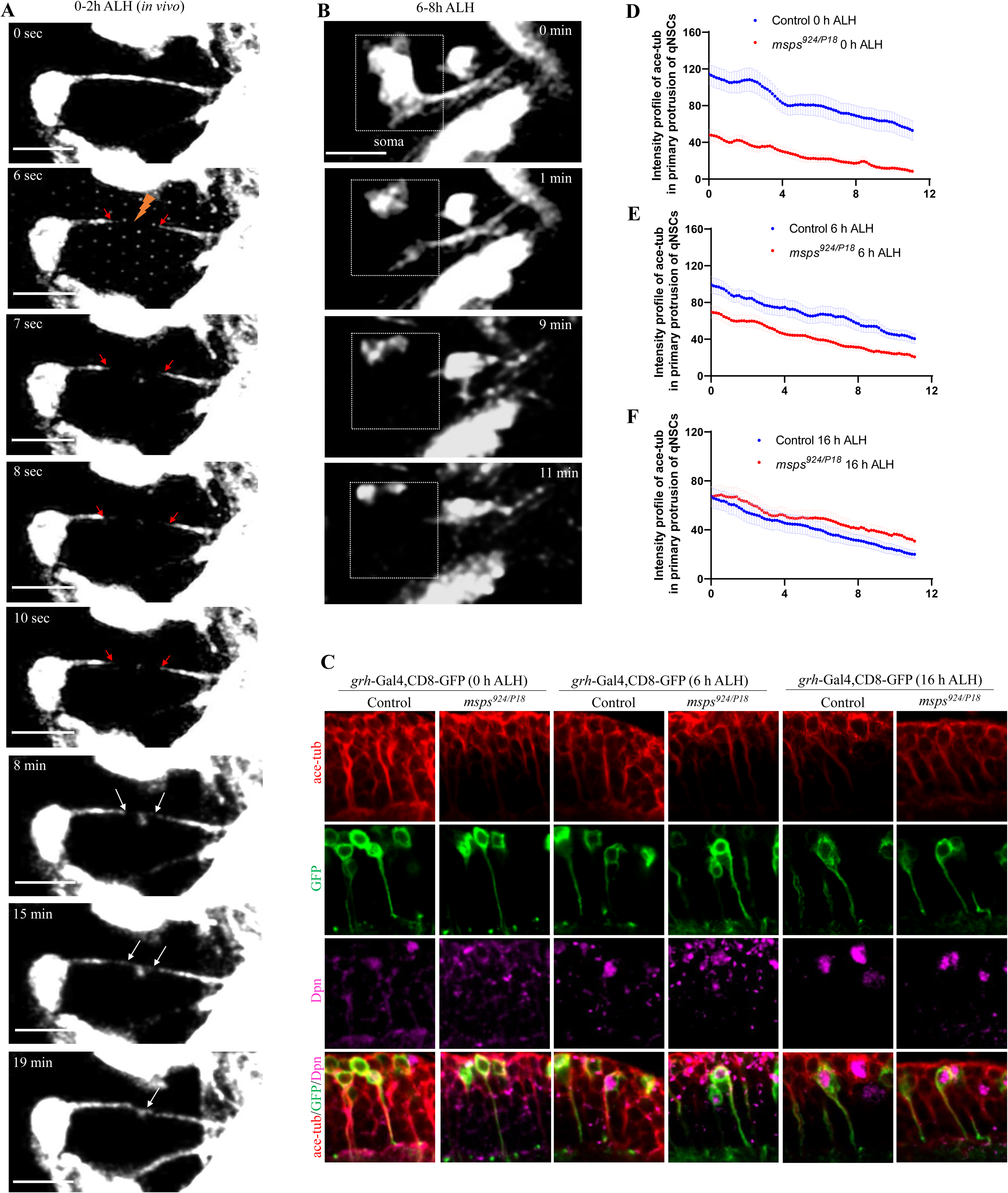
Quiescent NSC protrusions can regenerate upon injury *in vivo*. A) Time series of a qNSC in the larval brain from live whole larva (*in vivo*) at 0-2h ALH from *grh*-Gal4; *UAS*-mCD8-GFP ablated in the middle region of the protrusion (lightning bolt). After ablation primary protrusion of qNSCs show slight re-coil suggesting injury and not bleaching (red arrows). Complete regeneration is observed by 19 minutes (white arrows). 100% of the qNSCs from *in vivo* larval brains ablated at 0-2h ALH were capable of complete regeneration, n=4. B) Time series of a qNSC in the larval brain from live whole larva (*in vivo*) at 6-8h ALH from *grh*-Gal4; *UAS*-mCD8-GFP ablated at the cell body (dotted box). Ablation at the cell body resulted in the death of 100% of qNSCs, n=5. C) The larval ventral nerve cord (VNC) at 0h, 6h and 16h ALH, including the control (*UAS*-mCD8-GFP) and *msps^924/P18^* (trans-heterozygous *msps^924^/msps^P18^*) driven by *grh*-Gal4 were labelled with Ace-tub, Dpn, and GFP. D) Quantification of Ace-tub intensity in the primary protrusion of qNSCs in the control (n=28) and *msps^924/P18^*(n=21) VNC at 0h ALH. E) Quantification of Ace-tub intensity in the primary protrusion of qNSCs in the control (n=35) and *msps^924/P18^*(n=37) VNC at 6h ALH. F) Quantification of Ace-tub intensity in the primary protrusion of quiescent NSCs in the control (n=28) and *msps^924/P18^* (n=27) VNC at 16h ALH. Scale bars: 10 μm.

**Figure S2.**
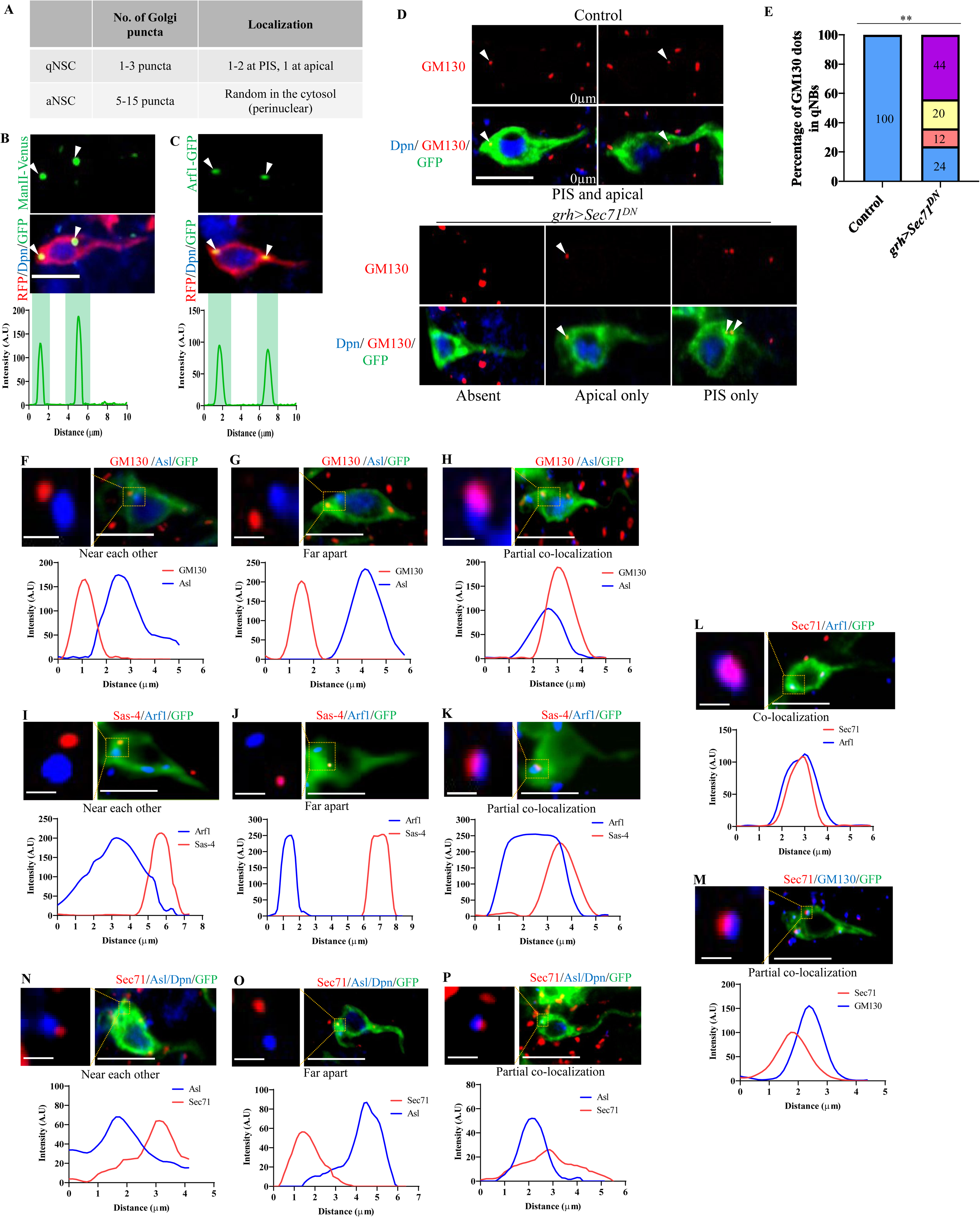
The Golgi apparatus localizes to the protrusion initial segment in *Drosophila* qNSCs. A) A table indicating the localization pattern of Golgi in quiescent and dividing NSCs. B) qNSCs in the larval brain expressing *UAS*-*ManII*-Venus driven by *grh*>CD8-RFP at 6h ALH were labelled with Dpn, GFP, and RFP. Arrowheads indicate *ManII*-Venus puncta localization at the apical and PIS regions=100%, n=42. Intensity graph showing the localization of *ManII*-Venus in the apical and PIS regions of qNSCs. C) qNSCs in the larval brain expressing *UAS*-Arf1-GFP driven by *grh*>CD8-RFP at 6h ALH were labelled with Dpn, GFP, and RFP. Arrowheads indicate Arf1-GFP puncta localization at the apical and PIS regions=100%, n=51. Intensity graph showing the localization of Arf1-GFP in the apical and PIS regions of qNSCs. D) qNSC protrusions in the larval brain at 6h ALH from control (*grh*-Gal4/*UAS-β-Gal* RNAi) and *grh*-Gal4/*UAS*-*Sec71^DN^* expressing *UAS*-CD8-GFP were labelled with antibodies against GM130, Dpn and GFP. Arrowheads indicate GM130 puncta. E) Quantification graph of the percentage of GM130 dots present per qNSC for genotypes in (D). GM130 puncta localization patterns: control, apical + PIS=100% (n=24); *UAS*-*Sec71^DN^*, apical + PIS=24%, apical only =12%, PIS only =20%, absent =44% (n=26). **p<0.01. F-H) qNSCs in the larval brain at 6h ALH under the control of *grh*-Gal4; *UAS*-CD8-GFP were labelled with GM130, Dpn, Asl and GFP. (F) Confocal micrograph and intensity graph showing the localization of GM130 and Asl near each other in qNSCs, 63.9%, n=36. (G) Confocal micrograph and intensity graph showing the localization of GM130 and Asl far apart from each other in qNSCs, 22.2%, n=36. (H) Confocal micrograph and intensity graph showing the partial colocalization of GM130 and Asl in qNSCs, 13.9%, n=36. I-K) qNSCs in the larval brain at 6h ALH under the control of *grh*-Gal4; *UAS*-CD8-GFP were labelled with Sas-4, Arf1 and GFP. (I) Confocal micrograph and intensity graph showing the localization of Arf1 and Sas-4 near each other in qNSCs, 68.4%, n=38. (J) Confocal micrograph and intensity graphs showing the localization of Arf1 and Sas-4 far apart from each other in qNSCs, 18.4%, n=38. (K) Intensity graphs showing the partial colocalization of Arf1 and Sas-4 in qNSCs, 13.2%, n=38. L) qNSCs in the larval brain at 6h ALH from *grh*-Gal4; *UAS*-CD8-GFP were labelled with antibodies against Sec71, Arf1 and GFP. Intensity graph showing the co-localization of Sec71 and Arf1 in qNSCs (100%, n=30). M) qNSCs in the larval brain at 6h ALH from *grh*-Gal4; *UAS*-CD8-GFP was labelled with antibodies against Sec71, GM130 and GFP. Intensity graph showing the partial co-localization of Sec71 and Arf1 in qNSCs (100%, n=33). N-P) qNSCs in the larval brain at 6h ALH from *grh*-Gal4; *UAS*-CD8-GFP were labelled with antibodies against Asl, Dpn, Sec71 and GFP. (N) Intensity graph showing the localization of Sec71 and Asl near each other in qNSCs, 69.2%, n=26. (O) Intensity graph showing the localization of Sec71 and Asl far apart from each other in qNSCs, 15.4%, n=26. (P) Intensity graph showing the partial co-localization of Sec71 and Asl in qNSCs, 15.4%, n=26. Scale bars: 1μm and 5μm.

**Figure S3.**
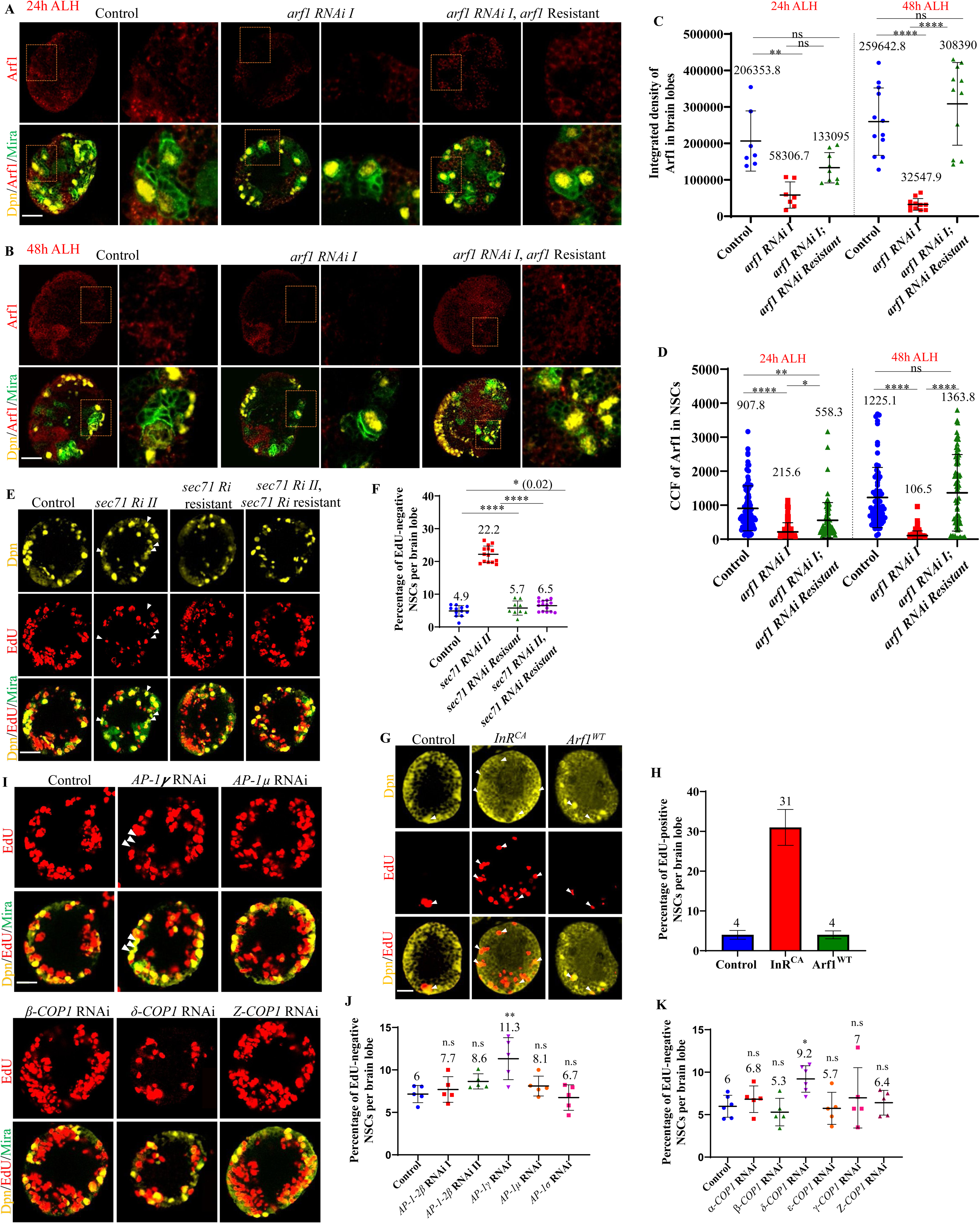
The Golgi proteins Arf1 and its GEF Sec71 are required for qNSC reactivation. A) Larval brains at 24h ALH, including the control (*grh*-Gal4 /*UAS-β-Gal* RNAi), *UAS*-*arf1 RNAi* I, and *UAS*-*arf1 RNAi* I, *UAS*-*arf1 RNAi* I resistant, driven by *grh*-Gal4 were labelled for Arf1, Dpn, and Mira. The boxed region on the left is magnified in the right panel. B) Larval brains at 48h ALH, including the control (*grh*-Gal4 /*UAS-β-Gal* RNAi), *UAS*-*arf1 RNAi* I, and *UAS*-*arf1 RNAi I*, *UAS*-*arf1 RNAi* I resistant, driven by *grh*-Gal4 were labelled for Arf1, Dpn, and Mira. The boxed region on the left is magnified in the right panel. C) Quantification graph of Arf1 intensity in whole-brain lobes from genotypes in (A and B). At 24h ALH, control, 206354 A.U., n=7 BL; *UAS*-*arf1 RNAi I*, 58307 A.U., n=7 BL; *UAS*-*arf1 RNAi I*, *UAS*-*arf1 RNAi I* resistant, 133095 A.U., n=9 BL. At 48h ALH, control, 259643 A.U., n=12 BL; *UAS*-*arf1 RNAi I*, 32548 A.U., n=12 BL; *UAS*-*arf1 RNAi I*, *UAS*-*arf1 RNAi I* resistant, 308390 A.U., n=11 BL. **p<0.01, ****p<0.0001, ns-non-significant. D) Quantification graph of the corrected total cell fluorescence (CCF) of Arf1 in NSCs for genotypes in (A and B). CCF = Integrated Density – (area of selected cell × mean fluorescence of background readings). At 24h ALH, control, 907.8 A.U., n=93; *UAS*-*arf1 RNAi I*, 215.6 A.U., n=70; *UAS*-*arf1 RNAi I*, *UAS*-*arf1 RNAi I* resistant, 558.3 A.U., n=83. At 48h ALH, control, 1225.1 A.U., n=71; *UAS*-*arf1 RNAi I*, 106.5 A.U., n=109; *UAS*-*arf1 RNAi I*, *UAS*-*arf1 RNAi I* resistant, 1363.8 A.U., n=71. *p<0.05, **p<0.01, ****p<0.0001, ns- non-significant. E) Larval brains at 24h ALH, including the control (*grh*-Gal4 /*UAS*-*β-Gal* RNAi), *UAS*-*sec71 RNAi II*, *UAS*-*sec71RNAi II* resistant, and *UAS*-*sec71 RNAi II*, *UAS*-*sec71RNAi II* resistant, driven by *grh*-Gal4 were analyzed for EdU incorporation. NSCs were marked by Dpn and Mira. Arrowheads indicate EdU-negative NSCs. F) Quantification graph of EdU-negative NSCs per brain lobe for genotypes in (E). Control, 4.9%, n=12 BL; *sec71 RNAi* II, 22.2% n=14 BL; *sec71* RNAi II resistant, 5.7%, n=9 BL; *sec71* RNAi II, *sec71* RNAi II resistant 6.5%, n=15 BL. ****p<0.0001, *p<0.05. G) Larval brains at 24h ALH, including the control (*grh*-Gal4 /*UAS-β-Gal* RNAi), positive control (*grh*-Gal4 /*UAS*-*InR^AD^*) and *UAS*-*Arf1^WT^*, driven by *grh*-Gal4 were analyzed for EdU incorporation under nutritional restriction (NR) conditions. Arrowheads indicate EdU-positive NSCs. H) Quantification graph of EdU-positive NSCs per brain lobe for genotypes in (G). Control, 4%, n=13 BL; *UAS*-*InR^AD^*, 31%, n=16; *UAS*-*Arf1^WT^*, 4%, n=14 BL. ****p<0.0001, ns- non-significant. I) Larval brains at 24h ALH, including the control (*UAS-β-Gal* RNAi), various RNAi against AP-1 and COP1 subunits, driven by *grh*-Gal4; *UAS Dcr2* were analyzed for EdU incorporation. NSCs were marked by Dpn and Mira. Arrowheads indicate EdU-negative NSCs. J) Quantification graph of EdU-negative NSCs per brain lobe for control and various AP-1 subunit RNAi lines in (I). **p<0.01, ns- non-significant. K) Quantification graph of EdU-negative NSCs per brain lobe for control and various COP1 subunit RNAi lines in (I). *p<0.05, ns- non-significant. Scale bars: 10 μm.

**Figure S4.**
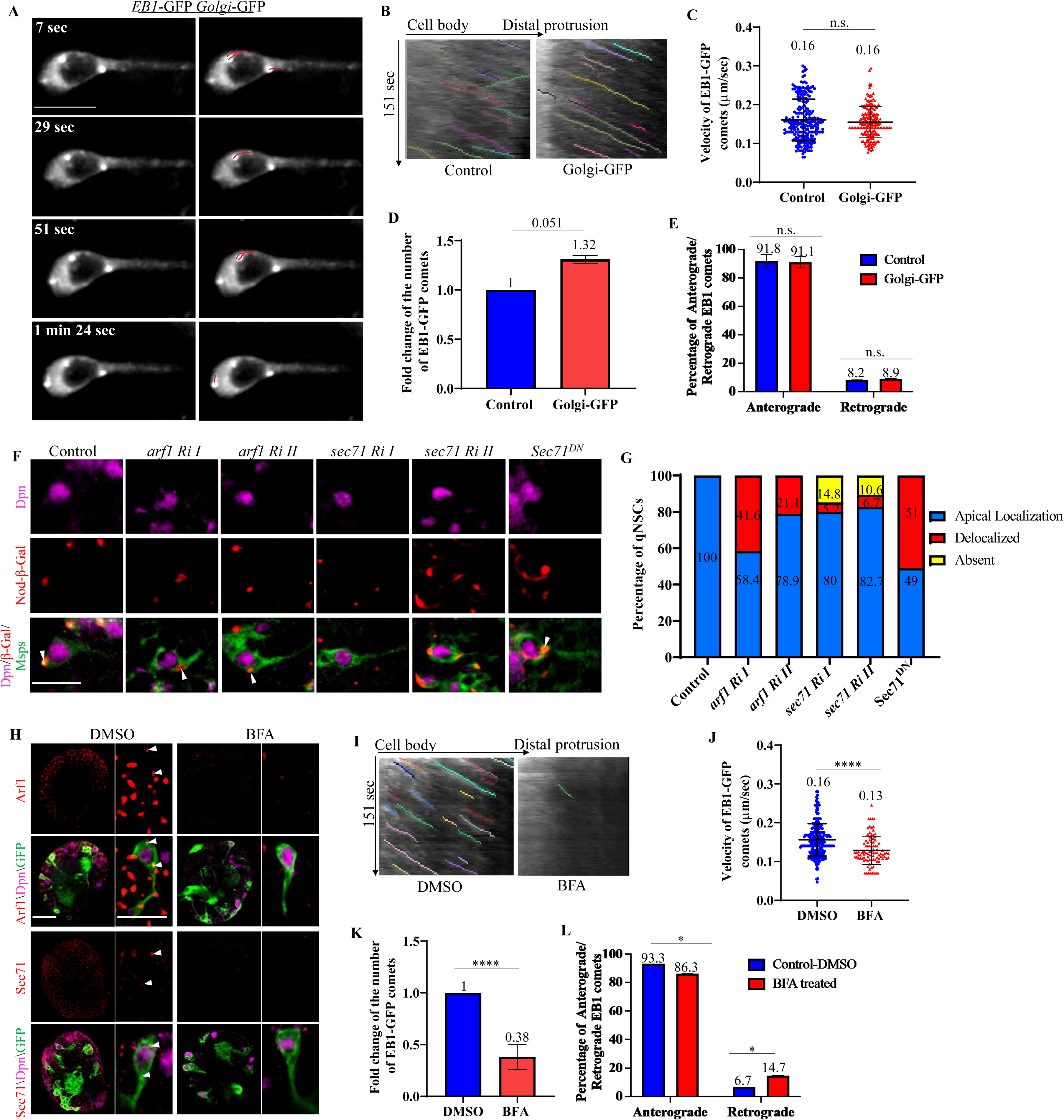
Golgi proteins Arf1 and Sec71 regulate microtubule assembly in the primary protrusions of qNSCs. A) Time series of a quiescent NSC from the larval brain at 6h ALH expressing *UAS*-EB1-GFP and *UAS*-*Golgi-GFP* driven by *grh*-Gal4. Red lines indicate the movement of EB1 comets emerging from Golgi present at the apical, lateral, and PIS regions. B) Kymograph of EB1-GFP comets movement in the primary protrusions of qNSCs in control (*grh*-Gal4/*UAS-β-Gal* RNAi) and *UAS*-*Golgi-*GFP (in A). C) Quantification graph of the velocity of EB1-GFP comets in the primary protrusion of qNSCs at 6h ALH from *Golgi*-GFP compared with control in (B). Control, 0.16 µm/second, n=207 comets, 32 qNSCs; *Golgi-*GFP, 0.16 µm/second, n=191 comets, 19 qNSCs. ns- non-significant. D) Quantification graph of fold changes in the number of EB1-GFP comets in the primary protrusion of qNSCs at 6h ALH in control and Golgi-GFP overexpression conditions in (B). Control, 1, n=207 comets; *Golgi-*GFP, 1.32, n=191 comets. ns- non-significant. E) Quantification graph of the percentage of anterograde and retrograde movements of EB1-GFP comets in the primary protrusion of qNSCs from *Golgi*-GFP compared with control in (B). Anterograde movements: control, 91.8%, n=248 comets; *Golgi-*GFP, 91.1%, n=174 comets. Retrograde movements: control, 8.2%, n=22 comets; *Golgi-*GFP, 8.9%, n=17 comets. ns- non-significant. F) Larval brains at 16h ALH, including the control (*UAS-β-Gal* RNAi), *arf1* RNAi I (*UAS*-*arf1* RNAi I /VDRC#23082GD), *arf1* RNAi II (*UAS*-*arf1* RNAi II /VDRC#103572KK), *sec71* RNAi I (*UAS*-*sec71* RNAi I/VDRC#100300KK), *sec71* RNAi II (*UAS*-*sec71* RNAi II/BSDC#32366) and *UAS*-*Sec71^DN^*with Nod-β-Gal expressed under *insc*-Gal4, *UAS*-Dcr2 and *insc*-Gal4 were labelled with β-Gal, Dpn, and Msps. Quiescent NSCs at the central brain (CB) are shown. G) Quantification graph of Nod-β-Gal localization in qNSCs from genotypes in (F). De-localization of Nod-β-Gal from the apical region: control, 0%, n=80; *arf1 RNAi I*, 41.6%, n=89, and *arf1 RNAi II*, 21.1%, n=71; *sec71 RNAi I*, 5.2%, n=115, and *sec71 RNAi II*, 6.7%, n=69; Sec71^DN^, 51%, n=55. Nod-β-Gal absent in qNSCs, control, *arf1 RNAi I*, and *arf1 RNAi II*= 0%; *sec71 RNAi I*, 14.8%, n=115, and *sec71 RNAi II*, 10.6%, n=69. ***p<0.001, *p<0.05, ns- non-significant. H) Larval brains at 6h ALH, including the control (DMSO-treated, n=9 BL) and BFA-treated brain lobes expressing *grh*-Gal4; *UAS*-CD8-GFP were stained for Arf1 or Sec71 (n=10 BL), Dpn and GFP. I) Kymograph of EB1-GFP comets movement in the primary protrusion of qNSCs from control (DMSO-treated) and BFA-treated larval brain lobes at 6h ALH in (H). J) Quantification graph of the velocity of EB1-GFP comets in the primary protrusion of qNSCs 6h ALH from various genotypes in (I). Control, velocity=0.16 μm/sec, n=241 comets in 20 qNSCs; BFA-treated, velocity=0.13 μm/sec, n=95 comets in 23 qNSCs. ****p<0.0001. K) Quantification graph of fold changes in the number of EB1-GFP comets in the primary protrusion of qNSCs 6h ALH upon BFA-treatment compared with the control in (I). Control, 1, n=241 comets in 20 qNSCs; BFA-treated, 0.38fold, n=95 comets in 23 qNSCs. ****p<0.0001. L) Quantification graph of the percentage of anterograde and retrograde movements of EB1-GFP comets in the primary protrusion of qNSCs upon BFA-treatment compared with control in (I). Anterograde movements in control, 93.3%, n=217; BFA-treated, 85.3%, n=81. Retrograde movements in control, 6.7%, n=24; BFA-treated, 14.7%, n=14. *p<0.05. Scale bars: 5μm and 10μm.

**Figure S5.**
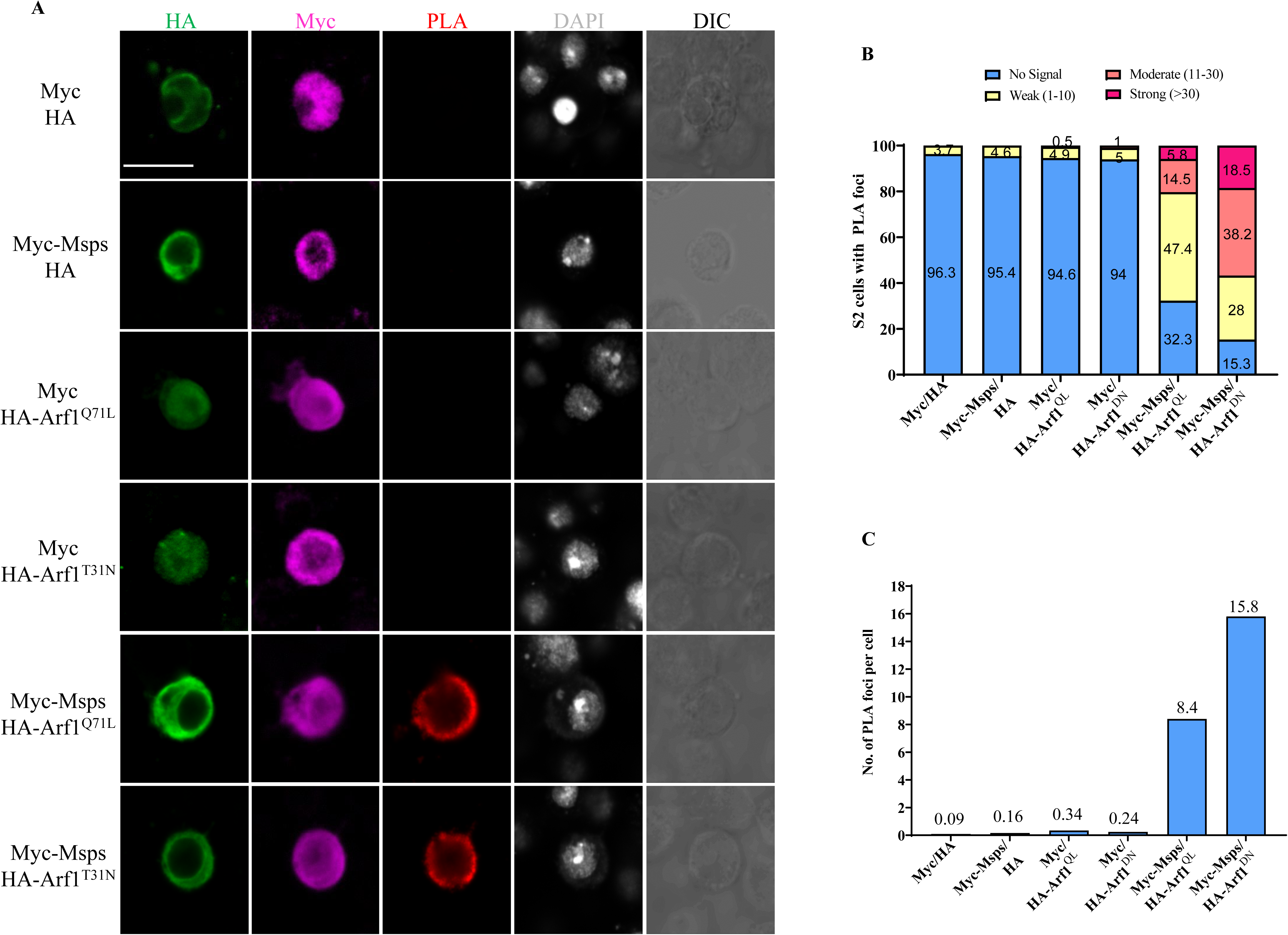
Arf1 and Msps physically associate in PLA assay. (A) *In situ* PLA assay among Arf1^Q71L^, Arf1^T31N^ and Msps in S2 cells. S2 cells transfected with two of the indicated plasmids (Myc, HA, Myc-Msps, HA-Arf1^Q71L^ and HA-Arf1^T31N^) were stained for HA, Myc, and DNA and detected for PLA signal. Cell outlines were determined by differential interference contrast (DIC) images. (B) Quantification graph showing the percentage of S2 cells with no PLA signal, or weak (1-10 foci), moderate (11–30 foci), or strong (>30 foci) PLA signals for (A). Myc and HA, no signal=96.3%, weak signal=3.7%, n=135; Myc-Msps and HA, no signal=95.4%, weak=4.6%, n=173; Myc and HA-Arf1^Q71L^, no signal=94.6%, weak=4.9%, moderate=0.5%, n=205; Myc and HA-Arf1^T31N^, no signal=94%, weak=5%, moderate=1%, n=199; Myc-Msps and HA-Arf1^Q71L^, no signal=32.3%, weak=47.4%, moderate=14.5%, strong=5.8% n=173; Myc-Msps and HA-Arf1^T31N^, no signal=15.3%, weak=28%, moderate=38.2%, strong=18.5%, n=157. (C) Quantification graph of the average number of PLA foci per cell in (A). Myc and HA, 0.09, n=135; HA and Myc-Msps, 0.16, n=173; Myc and HA-Arf1^Q71L^, 0.34, n=205; Myc and HA-Arf1^DN^ (Arf1^T31N^), 0.24, n=199; Myc-Msps and HA-Arf1^Q71L^, 8.4, n=173; Myc-Msps and HA-Arf1^T31N^, 15.8, n=157. Scale bars: 5 μm

**Figure S6.**
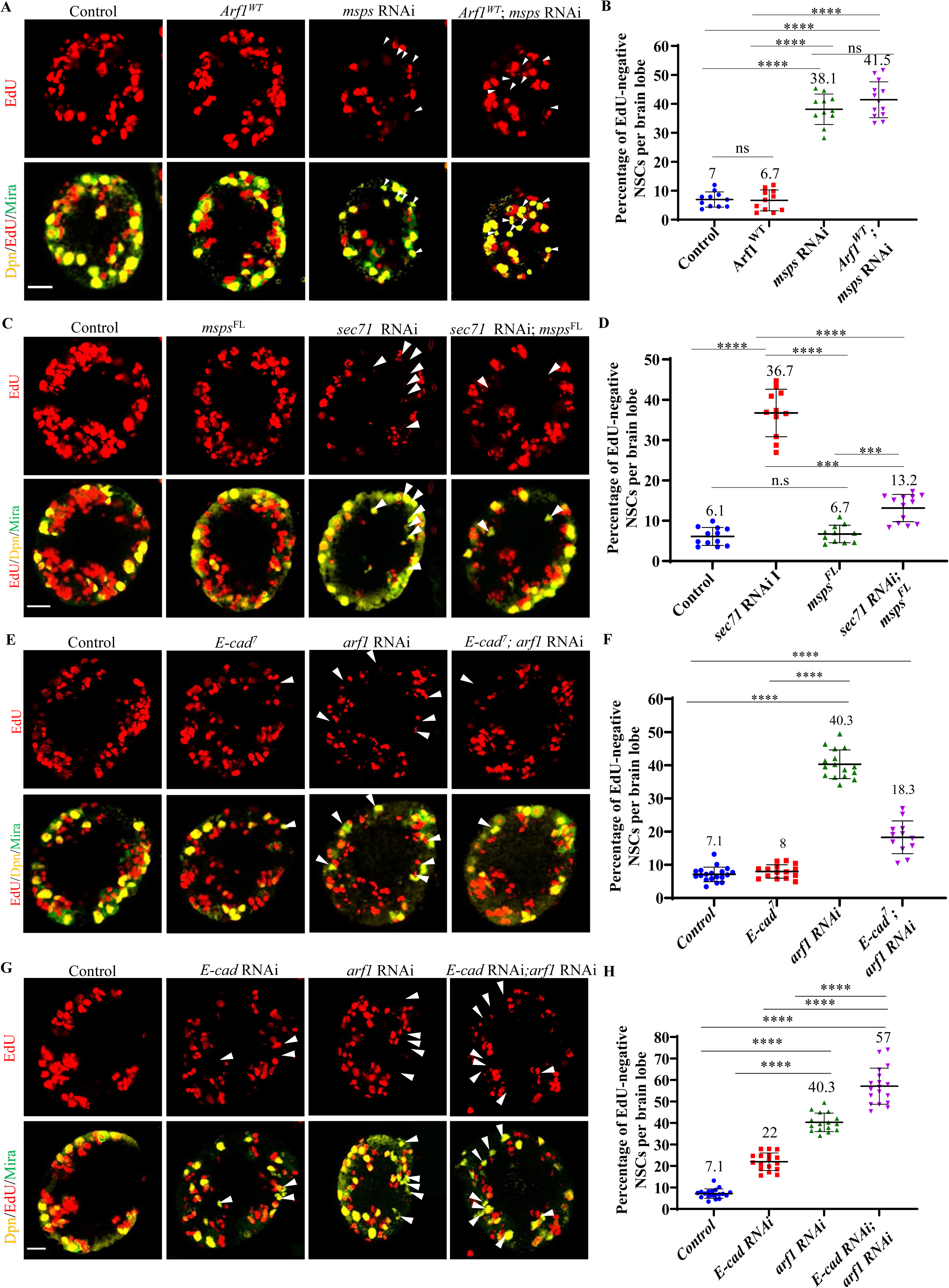
The microtubule regulator Msps and cell adhesion molecule E-cad act downstream of Arf1 to promote NSC reactivation. A) Larval brains at 24h ALH, including the control (*grh*-Gal4; *UAS*-Dcr2/*UAS-β-Gal* RNAi; *β-Gal* RNAi), *UAS*-*Arf1^WT^*; *β-Gal* RNAi, *UAS*-*msps* RNAi; *β-Gal* RNAi (VDRC#21982GD) and *UAS*-*Arf1^WT^*; *UAS*-*msps* RNAi, were analysed for EdU incorporation. NSCs were marked by Dpn and Mira. B) Quantification graph of EdU-negative NSCs per brain lobe for genotypes in (A). Control, 7%, n=11 BL; *UAS*-*Arf1^WT^*; *β-Gal* RNAi, 6.7%, n=11 BL; *UAS*-*msps* RNAi; *β-Gal* RNAi, 38.1, n=11 and *UAS*-*Arf1^WT^*; *UAS*-*msps* RNAi, 41.4%, n=13 BL. ****p<0.0001, ns-non-significant. C) Larval brains at 24h ALH from various genotypes were analysed for EdU incorporation. Control: *grh*-Gal4; *UAS*-Dcr2/*UAS-β-Gal* RNAi; *UAS-β-Gal* RNAi. *msps^FL^*: *grh*-Gal4 *UAS*-*msps^FL^*; *β-Gal* RNAi. *sec71* RNAi: *grh*-Gal4 *UAS*-*sec71* RNAi I (VDRC#100300KK); *β-Gal* RNAi*. sec71* RNAi *msps^FL^*: *grh*-Gal4 *UAS*-*sec71* RNAi I; *UAS*-*msps^FL^.* NSCs were marked by Dpn and Mira. White arrowheads point to NSCs without EdU incorporation. D) Quantification graph of EdU-negative NSCs per brain lobe for genotypes in (C). Control, 6.1%, n=12 BL; *sec71* RNAi I, 36.7%, n=11; *msps^FL^*, 6.7%, n=11 BL; and *sec71* RNAi; *msps^FL^*, 13.2%, n=13 BL. ****p<0.0001, ***p<0.001, ns- non-significant. E) Larval brains at 24h ALH, including the control (*grh*-Gal4, *UAS*-*mCD8-GFP*; *UAS*-Dcr2/*UAS-β-Gal* RNAi), *UAS*-*E-cad^7^*, *UAS*-*arf1* RNAi I (VDRC#23082GD) and *UAS*-*E-cad^7^*; *UAS*-*arf1* RNAi I, were analysed for EdU incorporation. NSCs were marked by Dpn and Mira. White arrowheads point to NSCs without EdU incorporation. F) Quantification graph of EdU-negative NSCs per brain lobe for genotypes in (E). Control, 7.1%, n=18 BL; *UAS*-*E-cad^7^*, 8%, n=15 BL; *UAS*-*arf1* RNAi I, 40.3%, n=16 and *UAS*-*E-cad^7^*; *UAS*-*arf1* RNAi I, 18.3%, n=12 BL. ****p<0.0001. G) Larval brains at 24h ALH, including the control (*grh*-Gal4, *UAS mCD8-GFP*; *UAS*-Dcr2/*UAS-β-Gal* RNAi), *UAS*-*E-cad* RNAi I (BDSC#38207), *UAS*-*arf1* RNAi I (VDRC#23082GD) and *UAS*-*E-cad* RNAi; *UAS*-*arf1* RNAi I, were analysed for EdU incorporation. NSCs were marked by Dpn and Mira. H) Quantification graph of EdU-negative NSCs per brain lobe for genotypes in (G). Control, 7.1%, n=18 BL; *UAS*-*E-cad* RNAi I, 22%, n=16 BL; *UAS*-*arf1* RNAi I, 40.3%, n=16 and *UAS*-*E-cad* RNAi; *UAS*-*arf1* RNAi I, 57%, n=18 BL. ****p<0.0001. White arrowheads point to NSCs without EdU incorporation. Scale bars: 10 μm

## References

1. Doetsch, F., Caille, I., Lim, D. A., Garcia-Verdugo, J. M. & Alvarez-Buylla, A. Subventricular zone astrocytes are neural stem cells in the adult mammalian brain. Cell 97, 703–716 (1999). https://doi.org:10.1016/s0092-8674(00)80783-7

2. Morshead, C. M. et al. Neural stem cells in the adult mammalian forebrain: a relatively quiescent subpopulation of subependymal cells. Neuron 13, 1071–1082 (1994). https://doi.org:10.1016/0896-6273(94)90046-9

3. Fabel, K. & Kempermann, G. Physical activity and the regulation of neurogenesis in the adult and aging brain. Neuromolecular Med 10, 59–66 (2008). https://doi.org:10.1007/s12017-008-8031-4

4. Lugert, S. et al. Quiescent and active hippocampal neural stem cells with distinct morphologies respond selectively to physiological and pathological stimuli and aging. Cell Stem Cell 6, 445–456 (2010). https://doi.org:10.1016/j.stem.2010.03.017

5. Baser, A., Skabkin, M. & Martin-Villalba, A. Neural Stem Cell Activation and the Role of Protein Synthesis. Brain Plast 3, 27–41 (2017). https://doi.org:10.3233/BPL-160038

6. Cloetta, D. et al. Inactivation of mTORC1 in the developing brain causes microcephaly and affects gliogenesis. J Neurosci 33, 7799–7810 (2013). https://doi.org:10.1523/JNEUROSCI.3294-12.2013

7. Ding, W. Y., Huang, J. & Wang, H. Waking up quiescent neural stem cells: Molecular mechanisms and implications in neurodevelopmental disorders. PLoS Genet 16, e1008653 (2020). https://doi.org:10.1371/journal.pgen.1008653

8. Otsuki, L. & Brand, A. H. Quiescent Neural Stem Cells for Brain Repair and Regeneration: Lessons from Model Systems. Trends Neurosci 43, 213–226 (2020). https://doi.org:10.1016/j.tins.2020.02.002

9. Isshiki, T., Pearson, B., Holbrook, S. & Doe, C. Q. Drosophila neuroblasts sequentially express transcription factors which specify the temporal identity of their neuronal progeny. Cell 106, 511–521 (2001). https://doi.org:10.1016/s0092-8674(01)00465-2

10. Lai, S. L. & Doe, C. Q. Transient nuclear Prospero induces neural progenitor quiescence. Elife 3 (2014). https://doi.org:10.7554/eLife.03363

11. Tsuji, T., Hasegawa, E. & Isshiki, T. Neuroblast entry into quiescence is regulated intrinsically by the combined action of spatial Hox proteins and temporal identity factors. Development 135, 3859–3869 (2008). https://doi.org:10.1242/dev.025189

12. Andersen, R. O., Turnbull, D. W., Johnson, E. A. & Doe, C. Q. Sgt1 acts via an LKB1/AMPK pathway to establish cortical polarity in larval neuroblasts. Dev Biol 363, 258–265 (2012). https://doi.org:10.1016/j.ydbio.2011.12.047

13. Colombani, J. et al. A nutrient sensor mechanism controls Drosophila growth. Cell 114, 739–749 (2003). https://doi.org:10.1016/s0092-8674(03)00713-x

14. Britton, J. S. & Edgar, B. A. Environmental control of the cell cycle in Drosophila: nutrition activates mitotic and endoreplicative cells by distinct mechanisms. Development 125, 2149–2158 (1998). https://doi.org:10.1242/dev.125.11.2149

15. Chell, J. M. & Brand, A. H. Nutrition-responsive glia control exit of neural stem cells from quiescence. Cell 143, 1161–1173 (2010). https://doi.org:10.1016/j.cell.2010.12.007

16. Sousa-Nunes, R., Yee, L. L. & Gould, A. P. Fat cells reactivate quiescent neuroblasts via TOR and glial insulin relays in Drosophila. Nature 471, 508–512 (2011). https://doi.org:10.1038/nature09867

17. Mairet-Coello, G., Tury, A. & DiCicco-Bloom, E. Insulin-like growth factor-1 promotes G(1)/S cell cycle progression through bidirectional regulation of cyclins and cyclin-dependent kinase inhibitors via the phosphatidylinositol 3-kinase/Akt pathway in developing rat cerebral cortex. J Neurosci 29, 775–788 (2009). https://doi.org:10.1523/JNEUROSCI.1700-08.2009

18. Yan, Y. P., Sailor, K. A., Vemuganti, R. & Dempsey, R. J. Insulin-like growth factor-1 is an endogenous mediator of focal ischemia-induced neural progenitor proliferation. Eur J Neurosci 24, 45–54 (2006). https://doi.org:10.1111/j.1460-9568.2006.04872.x

19. Arsenijevic, Y., Weiss, S., Schneider, B. & Aebischer, P. Insulin-like growth factor-I is necessary for neural stem cell proliferation and demonstrates distinct actions of epidermal growth factor and fibroblast growth factor-2. J Neurosci 21, 7194–7202 (2001).

20. Juanes, M. et al. Three novel IGF1R mutations in microcephalic patients with prenatal and postnatal growth impairment. Clin Endocrinol (Oxf) 82, 704–711 (2015). https://doi.org:10.1111/cen.12555

21. Li, S. et al. An intrinsic mechanism controls reactivation of neural stem cells by spindle matrix proteins. Nat Commun 8, 122 (2017). https://doi.org:10.1038/s41467-017-00172-9

22. Ding, R., Weynans, K., Bossing, T., Barros, C. S. & Berger, C. The Hippo signalling pathway maintains quiescence in Drosophila neural stem cells. Nat Commun 7, 10510 (2016). https://doi.org:10.1038/ncomms10510

23. Poon, C. L., Mitchell, K. A., Kondo, S., Cheng, L. Y. & Harvey, K. F. The Hippo Pathway Regulates Neuroblasts and Brain Size in Drosophila melanogaster. Curr Biol 26, 1034–1042 (2016). https://doi.org:10.1016/j.cub.2016.02.009

24. Gil-Ranedo, J. et al. STRIPAK Members Orchestrate Hippo and Insulin Receptor Signaling to Promote Neural Stem Cell Reactivation. Cell Rep 27, 2921–2933 e2925 (2019). https://doi.org:10.1016/j.celrep.2019.05.023

25. Huang, J. & Wang, H. Hsp83/Hsp90 Physically Associates with Insulin Receptor to Promote Neural Stem Cell Reactivation. Stem Cell Reports 11, 883–896 (2018). https://doi.org:10.1016/j.stemcr.2018.08.014

26. Ly, P. T. et al. CRL4Mahj E3 ubiquitin ligase promotes neural stem cell reactivation. PLoS Biol 17, e3000276 (2019). https://doi.org:10.1371/journal.pbio.3000276

27. Otsuki, L. & Brand, A. H. Cell cycle heterogeneity directs the timing of neural stem cell activation from quiescence. Science 360, 99–102 (2018). https://doi.org:10.1126/science.aan8795

28. Stone, M. C., Roegiers, F. & Rolls, M. M. Microtubules have opposite orientation in axons and dendrites of Drosophila neurons. Mol Biol Cell 19, 4122–4129 (2008). https://doi.org:10.1091/mbc.E07-10-1079

29. Kapitein, L. C. & Hoogenraad, C. C. Which way to go? Cytoskeletal organization and polarized transport in neurons. Mol Cell Neurosci 46, 9–20 (2011). https://doi.org:10.1016/j.mcn.2010.08.015

30. Lu, W., Lakonishok, M. & Gelfand, V. I. Kinesin-1-powered microtubule sliding initiates axonal regeneration in Drosophila cultured neurons. Mol Biol Cell 26, 1296–1307 (2015). https://doi.org:10.1091/mbc.E14-10-1423

31. Bornens, M. The centrosome in cells and organisms. Science 335, 422–426 (2012). https://doi.org:10.1126/science.1209037

32. Conduit, P. T., Wainman, A. & Raff, J. W. Centrosome function and assembly in animal cells. Nat Rev Mol Cell Biol 16, 611–624 (2015). https://doi.org:10.1038/nrm4062

33. Deng, Q., Tan, Y. S., Chew, L. Y. & Wang, H. Msps governs acentrosomal microtubule assembly and reactivation of quiescent neural stem cells. EMBO J 40, e104549 (2021). https://doi.org:10.15252/embj.2020104549

34. Martin, M. & Akhmanova, A. Coming into Focus: Mechanisms of Microtubule Minus-End Organization. Trends Cell Biol 28, 574–588 (2018). https://doi.org:10.1016/j.tcb.2018.02.011

35. Rios, R. M. The centrosome-Golgi apparatus nexus. Philos Trans R Soc Lond B Biol Sci 369 (2014). https://doi.org:10.1098/rstb.2013.0462

36. Sanders, A. A. & Kaverina, I. Nucleation and Dynamics of Golgi-derived Microtubules. Front Neurosci 9, 431 (2015). https://doi.org:10.3389/fnins.2015.00431

37. Cherfils, J. Arf GTPases and their effectors: assembling multivalent membrane-binding platforms. Curr Opin Struct Biol 29, 67–76 (2014). https://doi.org:10.1016/j.sbi.2014.09.007

38. Ferland, R. J. et al. Disruption of neural progenitors along the ventricular and subventricular zones in periventricular heterotopia. Hum Mol Genet 18, 497–516 (2009). https://doi.org:10.1093/hmg/ddn377

39. Ge, X. et al. Missense-depleted regions in population exomes implicate ras superfamily nucleotide-binding protein alteration in patients with brain malformation. NPJ Genom Med 1 (2016). https://doi.org:10.1038/npjgenmed.2016.36

40. Lee, M. J., Gergely, F., Jeffers, K., Peak-Chew, S. Y. & Raff, J. W. Msps/XMAP215 interacts with the centrosomal protein D-TACC to regulate microtubule behaviour. Nat Cell Biol 3, 643–649 (2001). https://doi.org:10.1038/35083033

41. Mishra, B. et al. Using microfluidics chips for live imaging and study of injury responses in Drosophila larvae. J Vis Exp, e50998 (2014). https://doi.org:10.3791/50998

42. Chen, K. et al. Arl2-and Msps-dependent microtubule growth governs asymmetric division. J Cell Biol 212, 661–676 (2016). https://doi.org:10.1083/jcb.201503047

43. Ori-McKenney, K. M., Jan, L. Y. & Jan, Y. N. Golgi outposts shape dendrite morphology by functioning as sites of acentrosomal microtubule nucleation in neurons. Neuron 76, 921–930 (2012). https://doi.org:10.1016/j.neuron.2012.10.008

44. Kondylis, V. & Rabouille, C. The Golgi apparatus: lessons from Drosophila. FEBS Lett 583, 3827–3838 (2009). https://doi.org:10.1016/j.febslet.2009.09.048

45. Wei, J. H. & Seemann, J. Unraveling the Golgi ribbon. Traffic 11, 1391–1400 (2010). https://doi.org:10.1111/j.1600-0854.2010.01114.x

46. Gillingham, A. K. & Munro, S. The small G proteins of the Arf family and their regulators. Annu Rev Cell Dev Biol 23, 579–611 (2007). https://doi.org:10.1146/annurev.cellbio.23.090506.123209

47. Wang, Y. et al. Sec71 functions as a GEF for the small GTPase Arf1 to govern dendrite pruning of Drosophila sensory neurons. Development 144, 1851–1862 (2017). https://doi.org:10.1242/dev.146175

48. Donaldson, J. G. & Jackson, C. L. ARF family G proteins and their regulators: roles in membrane transport, development and disease. Nat Rev Mol Cell Biol 12, 362–375 (2011). https://doi.org:10.1038/nrm3117

49. Popoff, V., Adolf, F., Brugger, B. & Wieland, F. COPI budding within the Golgi stack. Cold Spring Harb Perspect Biol 3, a005231 (2011). https://doi.org:10.1101/cshperspect.a005231

50. Mukherjee, A., Brooks, P. S., Bernard, F., Guichet, A. & Conduit, P. T. Microtubules originate asymmetrically at the somatic golgi and are guided via Kinesin2 to maintain polarity within neurons. Elife 9 (2020). https://doi.org:10.7554/eLife.58943

51. Weiner, A. T. et al. Endosomal Wnt signaling proteins control microtubule nucleation in dendrites. PLoS Biol 18, e3000647 (2020). https://doi.org:10.1371/journal.pbio.3000647

52. Zhou, W. et al. GM130 is required for compartmental organization of dendritic golgi outposts. Curr Biol 24, 1227–1233 (2014). https://doi.org:10.1016/j.cub.2014.04.008

53. Niu, T. K., Pfeifer, A. C., Lippincott-Schwartz, J. & Jackson, C. L. Dynamics of GBF1, a Brefeldin A-sensitive Arf1 exchange factor at the Golgi. Mol Biol Cell 16, 1213–1222 (2005). https://doi.org:10.1091/mbc.e04-07-0599

54. Goodwin, S. S. & Vale, R. D. Patronin regulates the microtubule network by protecting microtubule minus ends. Cell 143, 263–274 (2010). https://doi.org:10.1016/j.cell.2010.09.022

55. Fredriksson, S. et al. Protein detection using proximity-dependent DNA ligation assays. Nat Biotechnol 20, 473–477 (2002). https://doi.org:10.1038/nbt0502-473

56. Gohl, C., Banovic, D., Grevelhorster, A. & Bogdan, S. WAVE forms hetero- and homo-oligomeric complexes at integrin junctions in Drosophila visualized by bimolecular fluorescence complementation. J Biol Chem 285, 40171–40179 (2010). https://doi.org:10.1074/jbc.M110.139337

57. Shyu, Y. J. & Hu, C. D. Fluorescence complementation: an emerging tool for biological research. Trends Biotechnol 26, 622–630 (2008). https://doi.org:10.1016/j.tibtech.2008.07.006

58. Lantz, V. A. & Miller, K. G. A class VI unconventional myosin is associated with a homologue of a microtubule-binding protein, cytoplasmic linker protein-170, in neurons and at the posterior pole of Drosophila embryos. J Cell Biol 140, 897–910 (1998). https://doi.org:10.1083/jcb.140.4.897

59. Goode, B. L. & Feinstein, S. C. Identification of a novel microtubule binding and assembly domain in the developmentally regulated inter-repeat region of tau. J Cell Biol 124, 769–782 (1994). https://doi.org:10.1083/jcb.124.5.769

60. Wu, Z. et al. Caenorhabditis elegans neuronal regeneration is influenced by life stage, ephrin signaling, and synaptic branching. Proc Natl Acad Sci U S A 104, 15132–15137 (2007). https://doi.org:10.1073/pnas.0707001104

61. Geoffroy, C. G., Hilton, B. J., Tetzlaff, W. & Zheng, B. Evidence for an Age-Dependent Decline in Axon Regeneration in the Adult Mammalian Central Nervous System. Cell Rep 15, 238–246 (2016). https://doi.org:10.1016/j.celrep.2016.03.028

62. Blanquie, O. & Bradke, F. Cytoskeleton dynamics in axon regeneration. Curr Opin Neurobiol 51, 60–69 (2018). https://doi.org:10.1016/j.conb.2018.02.024

63. Hellal, F. et al. Microtubule stabilization reduces scarring and causes axon regeneration after spinal cord injury. Science 331, 928–931 (2011). https://doi.org:10.1126/science.1201148

64. Sengottuvel, V. & Fischer, D. Facilitating axon regeneration in the injured CNS by microtubules stabilization. Commun Integr Biol 4, 391–393 (2011). https://doi.org:10.4161/cib.4.4.15552

65. De Camilli, P., Moretti, M., Donini, S. D., Walter, U. & Lohmann, S. M. Heterogeneous distribution of the cAMP receptor protein RII in the nervous system: evidence for its intracellular accumulation on microtubules, microtubule-organizing centers, and in the area of the Golgi complex. J Cell Biol 103, 189–203 (1986). https://doi.org:10.1083/jcb.103.1.189

66. Horton, A. C. et al. Polarized secretory trafficking directs cargo for asymmetric dendrite growth and morphogenesis. Neuron 48, 757–771 (2005). https://doi.org:10.1016/j.neuron.2005.11.005

67. Yang, S. Z. & Wildonger, J. Golgi Outposts Locally Regulate Microtubule Orientation in Neurons but Are Not Required for the Overall Polarity of the Dendritic Cytoskeleton. Genetics 215, 435–447 (2020). https://doi.org:10.1534/genetics.119.302979

68. Ruschel, J. et al. Axonal regeneration. Systemic administration of epothilone B promotes axon regeneration after spinal cord injury. Science 348, 347–352 (2015). https://doi.org:10.1126/science.aaa2958

69. Sengottuvel, V., Leibinger, M., Pfreimer, M., Andreadaki, A. & Fischer, D. Taxol facilitates axon regeneration in the mature CNS. J Neurosci 31, 2688–2699 (2011). https://doi.org:10.1523/JNEUROSCI.4885-10.2011

70. Erez, H. et al. Formation of microtubule-based traps controls the sorting and concentration of vesicles to restricted sites of regenerating neurons after axotomy. J Cell Biol 176, 497–507 (2007). https://doi.org:10.1083/jcb.200607098

71. Schmoranzer, J. & Simon, S. M. Role of microtubules in fusion of post-Golgi vesicles to the plasma membrane. Mol Biol Cell 14, 1558–1569 (2003). https://doi.org:10.1091/mbc.e02-08-0500

72. Cao, H. et al. Actin and Arf1-dependent recruitment of a cortactin-dynamin complex to the Golgi regulates post-Golgi transport. Nat Cell Biol 7, 483–492 (2005). https://doi.org:10.1038/ncb1246

73. Dubois, T. et al. Golgi-localized GAP for Cdc42 functions downstream of ARF1 to control Arp2/3 complex and F-actin dynamics. Nat Cell Biol 7, 353–364 (2005). https://doi.org:10.1038/ncb1244

74. Kjos, I., Vestre, K., Guadagno, N. A., Borg Distefano, M. & Progida, C. Rab and Arf proteins at the crossroad between membrane transport and cytoskeleton dynamics. Biochim Biophys Acta Mol Cell Res 1865, 1397–1409 (2018). https://doi.org:10.1016/j.bbamcr.2018.07.009

75. Koronakis, V. et al. WAVE regulatory complex activation by cooperating GTPases Arf and Rac1. Proc Natl Acad Sci U S A 108, 14449–14454 (2011). https://doi.org:10.1073/pnas.1107666108

76. Fucini, R. V., Chen, J. L., Sharma, C., Kessels, M. M. & Stamnes, M. Golgi vesicle proteins are linked to the assembly of an actin complex defined by mAbp1. Mol Biol Cell 13, 621–631 (2002). https://doi.org:10.1091/mbc.01-11-0547

77. Gurel, P. S., Hatch, A. L. & Higgs, H. N. Connecting the cytoskeleton to the endoplasmic reticulum and Golgi. Curr Biol 24, R660–R672 (2014). https://doi.org:10.1016/j.cub.2014.05.033

78. Bellouze, S. et al. Golgi fragmentation in pmn mice is due to a defective ARF1/TBCE cross-talk that coordinates COPI vesicle formation and tubulin polymerization. Hum Mol Genet 23, 5961–5975 (2014). https://doi.org:10.1093/hmg/ddu320

79. Hara, Y., Shagirov, M. & Toyama, Y. Cell Boundary Elongation by Non-autonomous Contractility in Cell Oscillation. Curr Biol 26, 2388–2396 (2016). https://doi.org:10.1016/j.cub.2016.07.003

80. Jakobs, M. A., Dimitracopoulos, A. & Franze, K. KymoButler, a deep learning software for automated kymograph analysis. Elife 8 (2019). https://doi.org:10.7554/eLife.42288

81. Goshima, G. & Vale, R. D. The roles of microtubule-based motor proteins in mitosis: comprehensive RNAi analysis in the Drosophila S2 cell line. J Cell Biol 162, 1003–1016 (2003). https://doi.org:10.1083/jcb.200303022

